# miR156-independent repression of the ageing pathway by longevity-promoting AHL proteins in Arabidopsis

**DOI:** 10.1101/2020.06.18.160234

**Authors:** Arezoo Rahimi, Omid Karami, Remko Offringa

**Affiliations:** Plant Developmental Genetics, Institute of Biology Leiden, Leiden University, Sylviusweg 72, 2333 BE Leiden, Netherlands

**Keywords:** Vegetative Phase Change, *AHL* genes, *SPL* genes, *miR156/miR157*, Arabidopsis

## Abstract

Plants age by transition through distinct developmental phases, with the juvenile- to adult vegetative and the adult vegetative to reproductive phase change as two consecutive important post-embryonic phase transitions. During the transition from the juvenile- to adult vegetative phase, also known as the vegetative phase change (VPC), many plants undergo specific morphological and physiological changes. In the model plant *Arabidopsis thaliana* (Arabidopsis), for example, the VPC is marked by clear heteroblastic changes in leaf shape and size and the appearance of trichomes on the abaxial side of leaves. The VPC and the vegetative to reproductive transition are promoted by the SQUAMOSA PROMOTOR BINDING PROTEIN-LIKE (SPL) family of transcription factors and repressed by miR156 and miR157 that target the *SPL* transcripts. Here we present data that the plant longevity promoting AT-HOOK MOTIF NUCLEAR LOCALIZED protein 15 (AHL15) and family members also repress the SPL-driven ageing pathway. Arabidopsis *ahl* loss-of-function mutants showed an accelerated VPC and flowering time, whereas *AHL15* overexpression dramatically delayed the VPC and flowering time in both Arabidopsis and *Nicotiana tabacum*. Expression analysis and tissue-specific *AHL15* overexpression revealed that *AHL15* affects the VPC and flowering time directly through its expression in the shoot apical meristem and young leaves. In addition, we found evidence that AHL15 represses *SPL* gene expression in a miR156/157-independent manner. The juvenile traits of *spl* loss-of-function mutants appeared to be dependent on the enhanced expression of the *AHL15* gene, providing evidence for a reciprocal negative feedback between *AHL15* and *SPL* genes. Antagonistically to AHLs, SPLs promote axillary meristem (AM) maturation and thus prevent vegetative growth from these meristems by repressing *AHL15* expression. Taken together, our results place AHL15 and family members at a central position in the SPL-driven ageing pathway as suppressors of the VPC, flowering time and AM maturation.

## Introduction

Plant development progresses through several distinct developmental phases, starting with embryogenesis and followed successively by the juvenile vegetative, adult vegetative, reproductive and gametophytic phase. In the juvenile vegetative phase, the plant is not competent to flower. Therefore, flowering requires the transition from juvenile to adult vegetative development, which is referred to as the vegetative phase change (VPC). The VPC generally leads to plant morphological changes such as increased internode length, adventitious root production and changes in leaf size and shape and trichome distribution, a situation also known as heteroblasty (Huijser and Schmid, 2011). In *Arabidopsis*, leaf heteroblasty provides a clear indicator of the VPC. Juvenile leaves have smooth margins, are rounder (length/width), and lack abaxial trichomes, whereas adult leaves have serrated margins, are more elongated (length/width ratio) and have abaxial trichomes (Telfer et al., 1997).

Compared to the adult-to-reproductive phase transition, which is one of the key-traits in crops, much less is known about the molecular mechanisms that mediate the juvenile-to-adult transition. However, recent progress in *Arabidopsis thaliana* (Arabidopsis) has demonstrated that microRNAs (miRNAs) miR156, miR157, and miR172, are major regulators of the juvenile-to-adult transition in Arabidopsis and other plant species (Poethig, 2013; Teotia and Tang, 2015). During the Arabidopsis life cycle the gradual decrease in *miR156/miR157* expression results in increased expression of the *SQUAMOSA-PROMOTER BINDING PROTEIN-LIKE (SPL) miR156/miR157* target genes. SPL transcription factors, in turn, promote the adult developmental program at the shoot apical meristem (SAM), resulting in the transition from juvenile to adult leaf production and eventually from vegetative to reproductive development (Wang et al., 2009; Wu et al., 2009; He et al., 2018). The gradual decrease of *miR156/miR157* expression during shoot maturation is accompanied by an SPL-induced gradual increase in *miR172* expression. miR172 promotes the development of trichomes on the abaxial side of leaves by repressing the expression of the APETALA2-LIKE (AP2-like) transcription factors TARGET OF EARLY ACTIVATION TAGGED1 (TOE1) and TOE2 (Wu et al., 2009). In addition, SPLs promote the other adult leaf traits such as leaf elongation and leaf serration independent of miR172. However, the genes that act downstream of SPLs and TOE1/TOE2 in juvenile-and-adult transition remain unidentified.

In eukaryotes, a wide range of DNA binding proteins have been identified that bind to the minor groove of DNA by a small AT-hook motif (Reeves, 2010). AT-hook proteins are considered as chromatin architectural factors involved in a diverse array of crucial cellular processes, including cell growth, -differentiation, -transformation, -proliferation, -death, and DNA replication and repair, by regulating chromatin remodelling, and gene transcription (Reeves, 2010; Sgarra et al., 2010; Ozturk et al., 2014). The Arabidopsis genome encodes 29 AT-Hook motif nuclear-Localized (AHL) proteins that containing either one or two AT-hook domains and a Plant and Prokaryote Conserved (PPC) domain (Fujimoto et al., 2004; Matsushita et al., 2007; Street et al., 2008; Zhao et al., 2013). These AHL proteins have been shown to be implicated in several aspects of plant growth and development, including hypocotyl growth (Street et al., 2008; Xiao et al., 2009; Zhao et al., 2013), root vascular tissue differentiation (Zhou et al., 2013), flower development (Ng et al., 2009), flowering time (Street et al., 2008; Xiao et al., 2009; Yun et al., 2012; Xu et al., 2013). Based on mutants and protein-protein interaction studies, the AHL family members have been proposed to bind AT-rich DNA regions as hetero-multimeric complexes. These AHL complexes use the AT-hook domains to anchor to AT-rich DNA regions and recruit other transcription factors through their interacting PPC domains (Zhao et al., 2013). In addition, it has been shown that AHL proteins repress transcription of several key developmental regulatory genes, possibly through modulation of the epigenetic code in the vicinity of their DNA binding regions (Lim et al., 2007; Ng et al., 2009; Yun et al., 2012). Some evidence has been obtained that AHL proteins function by altering the organization of the chromatin structure (Lim et al., 2007; Ng et al., 2009; Yun et al., 2012; Xu et al., 2013). However, since this plant-specific class of nuclear proteins has only been studied more recently, their exact mode of action is still elusive.

Recently, we have shown that the *AHL15* gene and other *AHL* family members are important for embryogenesis (Karami et al., 2020b) and that they promote plant longevity by delaying axillary meristem (AM) maturation (Karami et al., 2020a). In view of the seemingly antagonistic effect of *AHL15* and its family members on the plant ageing pathway, we here studied their possible role in the regulation of the VPC and flowering time. Our analyses in Arabidopsis and *Nicotiana tabacum* (tobacco) showed that *AHL15* overexpression prolongs the juvenile vegetative phase and delays flowering, whereas *ahl15* loss of function results in precocious appearance of adult vegetative traits. A more detailed analysis indicated that AHL15 delays developmental phase changes by repressing *SPL* gene expression in a miRNA-independent manner. We further show that in turn *AHL15* expression is repressed through feedback regulation by the SPLs.

## Results

### *AHL* genes delay the vegetative phase change and flowering time

When studying plants overexpressing *AHL15* (*p35S:AHL15*), we noticed that they initially produced much smaller rosette leaves than wild-type plants with a low number of adaxial trichomes (not shown), and that leaves produced three to four weeks later were similar in size to the first juvenile leaves of wild-type plants (Fig. 1A, B). More detailed analysis showed that *p35S:AHL15* plants produced leaves with a reduced length/width ratio (Fig. 1C) and had a significantly delayed VPC, as indicated by an increased number of leaves without abaxial trichomes, both under short day (SD) and long day (LD) conditions (Fig. 1D, E). By contrast, *ahl15* loss-of-function mutants developed leaves with a slightly increased length/width ratio with the VPC occurring 1 to 2 plastochrons earlier (Fig. 1A-E). Introduction of a *pAHL15:AHL15* genomic clone into the *ahl15* mutant background resulted again in wild-type leaf development (Fig. 1A-E), indicating that the phenotypes were caused by *ahl15* loss-of-function. *AHL15* is part of a large gene family in Arabidopsis, where it clusters together with two close homologs *AHL19* and *AHL20* (Karami et al., 2020a). In line with the previously reported high degree of functional redundancy between *AHL* genes (Street et al., 2008; Xiao et al., 2009; Zhao et al., 2013), *ahl15 ahl19 p35S:amiRAHL20* triple mutant plants showed a stronger increase in the leaf length/width ratio compared to the *ahl15* single or *ahl15 ahl19* double mutant. However, the timing of the VPC was comparable with that of the single or *ahl15 ahl19* double mutant plants under both SD and LD conditions (Fig. 1A-E).

**Figure 1.**
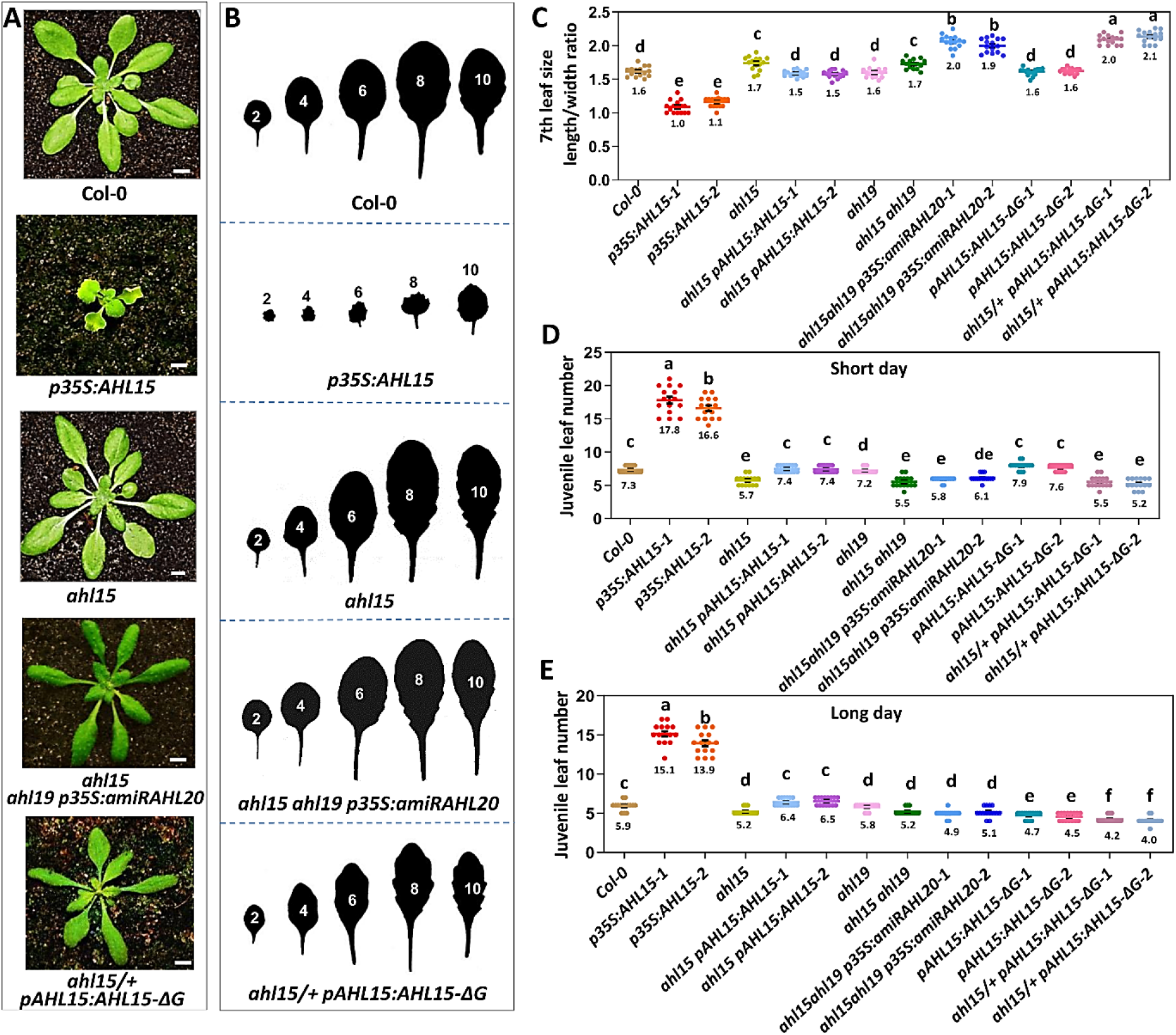
*AHL15* and family members delay the vegetative phase change in Arabidopsis. (**A**) The rosette phenotype of 5-week-old wild-type (Col-0), *p35S:AHL15*, *ahl15*, *ahl15 ahl19 p35S:amiRAHL20*, or *ahl15/+ pAHL15:AHL15-ΔG* plants grown under short-day (SD) conditions. Size bars indicate 1 cm. (**B**) Shape of the successive rosette leaves of 7-week-old of wild-type, *p35S:AHL15*, *ahl15 ahl19 p35S:amiRAHL20*, or *ahl15/+ pAHL15:AHL15-ΔG* plants grown under SD. (**C**) The length:width ratio of the 7 ^th^ leaf of 7-week-old wild-type, *p35S:AHL15*, *ahl15*, *ahl15 pAHL15:AHL15*, *ahl19*, *ahl15 ahl19*, *ahl15 ahl19 p35S:amiRAHL20*, *pAHL15:AHL15-ΔG* and *ahl15/+ pAHL15:AHL15-ΔG* plants grown under SD. (**D** and **E**) The juvenile leaf number (leaves without abaxial trichomes) in wild-type, *p35S:AHL15*, *ahl15*, *ahl15 pAHL15:AHL15*, *ahl19*, *ahl15 ahl19*, *ahl15 ahl19 p35S:amiRAHL20*, *pAHL15:AHL15-ΔG* and *ahl15/+ pAHL15:AHL15-ΔG* plants grown under SD (**D**) and under long day (**E**). Coloured dots in C-E indicate length:width ratio (C) and the number of leaves without abaxial trichomes (D,E) per plant (n = 15 biologically independent plants) per line, horizontal line and the number below this line indicate the mean and error bars indicate the s.e.m. **C-E** Different letters indicate statistically significant differences (P < 0.01) as determined by one-way ANOVA with Tukey’s honest significant difference post hoc test.

Previously, we have shown that expression of a mutant version of AHL15 lacking the conserved GRFEIL motif in the PPC domain under control of the *AHL15* promoter in the heterozygous mutant background (*ahl15+/−pAHL15:AHL15-ΔG*) leads to a dominant-negative effect that allows to overcome the functional redundancy between *AHL* genes (Karami et al., 2020a). Expression of *pAHL15:AHL15-ΔG* in the wild-type background only led to an acceleration of the VPC under LD conditions (Fig. 1A-E). However, ahl15*/+ pAHL15:AHL15-ΔG* plants showed the highest increase in leaf length/width ratio and also the strongest enhancement in timing of the VPC under LD conditions compared to wild type or to other *ahl* mutant lines (Fig. 1A-E). Since Arabidopsis plants require the VPC before they can shift to the reproductive phase, we also monitored the flowering time of our mutant lines. Our previous analysis already showed that *ahl15/+ pAHL15:AHL15-ΔG* plants flowered significantly earlier, as quantified by the number rosette leaves developed until flowering, under both LD and SD conditions compared to wild-type plants (Karami et al., 2020a). Further analysis showed that the single *ahl15* and *ahl19* loss-of-function mutations did not significantly affect flowering time, whereas *ahl15 ahl19 p35S:amiRAHL20* triple mutant plants showed a significant reduction in flowering time only under SD conditions (Supplementary Fig. 1A-C). In contrast, *p35S:AHL15* plants showed a significant delay of flowering under both SD and LD conditions (Supplementary Fig. 1A-C). These results indicate that *AHL* genes are important determinants of the VPC and, most likely as a result, also of the vegetative to reproductive phase transition (Supplementary Fig. 1A-C).

We also analysed the effect of *AHL15* overexpression on the same developmental phase changes in tobacco, a plant species from a different family, using available *p35S:AHL15-GR* tobacco lines (Karami et al., 2020a). In contrast to mock-treated *p35S:AHL15-GR* tobacco plants, which showed wild-type leaf development with three round juvenile leaves preceding the production of the typically larger and longer adult leaves, DEX-treated *p35S:AHL15-GR* plants formed many small, round juvenile leaves (Supplementary Fig. 2A). By spraying these plants once per week with DEX, the shoot apical meristem continued to produce small juvenile leaves, and plants could be kept in the juvenile vegetative state for more than a year, resulting in highly branched and bushy plants (Supplementary Fig. 2B and C). In contrast, *p35S:AHL15-GR* control plants that were not treated with DEX developed normally and flowered after two-month (not shown). This result indicates that *AHL15* overexpression can also strongly delay, if not prevent the VPC and flowering in a non-brassicaceae plant species, such as tobacco.

### *AHL15* delays the VPC through its expression in the SAM and in leaf primordia

To further understand the role of *AHL* genes in these developmental switches, we analysed the expression dynamics of *AHL15* and its close homologs *AHL19* and *AHL20* during the VPC. Quantitative RT-PCR (qPCR) experiments showed that expression of the three *AHL* genes was negatively correlated with shoot age, as expression of all three genes was higher in the shoot apex and young leaves of 1-or 2-week-old seedlings grown under short day (SD) conditions, and then significantly declined in these tissues in 3-week-old seedlings (Fig. 2A). This is in line with the timing of the VPC, which for Arabidopsis ecotype Col-0 has been reported to occur at 17-20 days after germination (DAG) under SD conditions (Xu et al., 2016). Similarly, analysis of *AHL15* expression in leaves and shoot apices using *pAHL15:GUS* reporter lines showed that *AHL15* was expressed throughout the shoot at 7 and 12 DAG (week 1 and week 2), that its expression declined in the shoot apical meristem (SAM) at 12 DAG, but that at 17 DAG (week 3) its expression was off or severely reduced in respectively the SAM or the new-formed leaves (Fig, 2B). The gene expression dynamics of *AHL15* and its close homologs supports their role in maintaining vegetative juvenile traits and thus suppressing the VPC.

**Figure 2.**
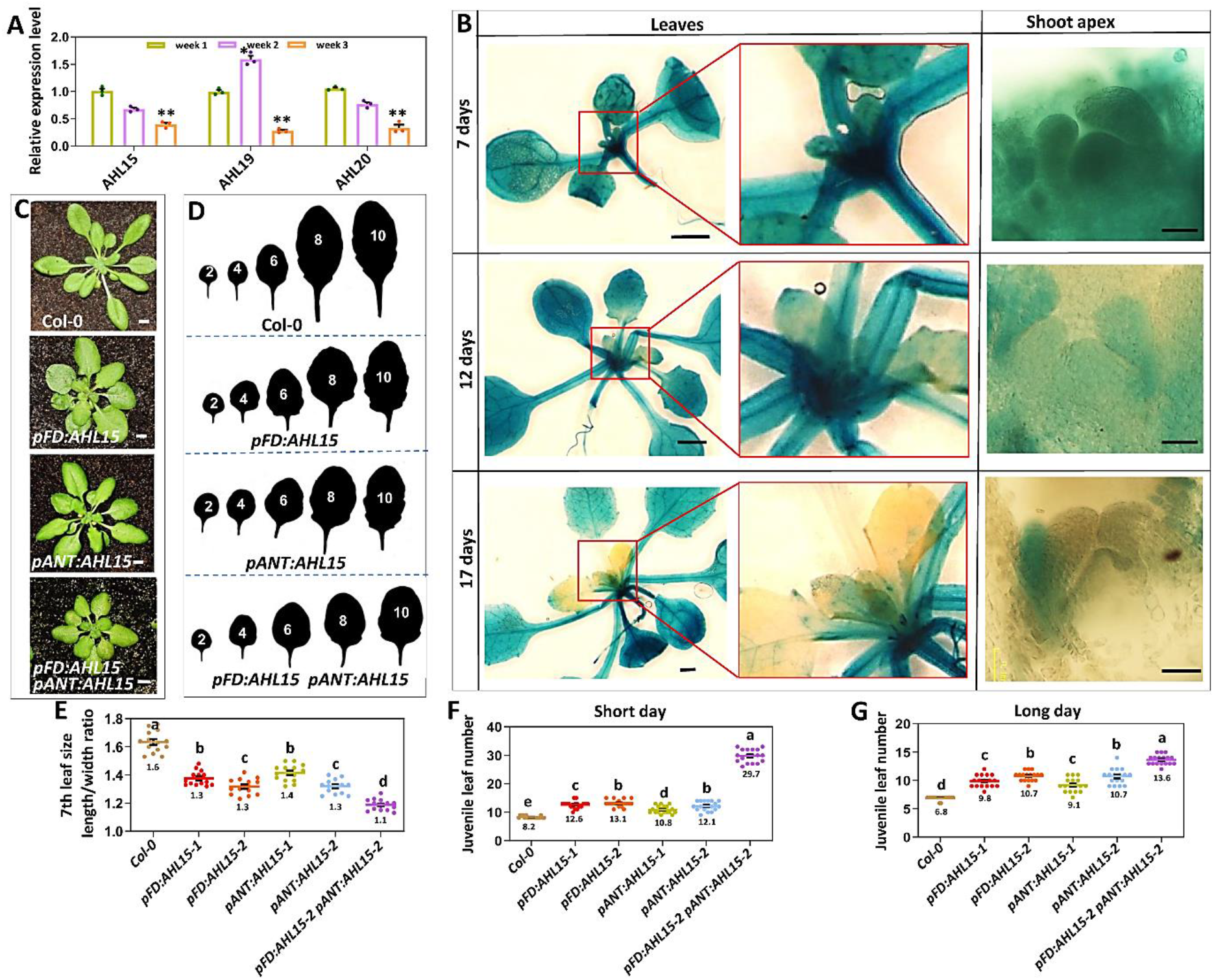
*AHL15* expression in the SAM and leaf primordia delays the VPC. (**A**) The relative expression level of *AHL15*, *AHL19*, and *AHL20* in the shoot apex and young leaves of 1, 2, and 3 week-old seedlings grown under SD conditions. Dots indicate the values of three biological replicates per plant line, bar indicates the mean, and error bars the s.e.m. The asterisk indicates a significant difference (*P< 0.05, **P< 0.01), as determined by a two-sided Student’s *t*-test. (**B**) Histochemical staining for GUS activity in 7-, 12-, and 17-day-old *pAHL15:GUS* seedlings. Images show overview of leaves (left two rows) and details shoot apex (right row). (**C**) The rosette phenotype of 5-week-old wild-type (Col-0), *pFD:AHL15*, *pANT:AHL15* and *pFD:AHL15 pANT:AHL15* plants grown under SD conditions. (**D**) Shape successive rosette leaves of 7-week-old of wild-type, *pFD:AHL15*, *pANT:AHL15* and *pFD:AHL15 pANT:AHL15* plants grown under SD. (**E**) The length:width ratio of the 7 th leaf of 7-week-old wild-type, *pFD:AHL15*, *pANT:AHL15* and *pFD:AHL15 pANT:AHL15* plants grown under SD. (**F** and **H**) The juvenile leaf number (leaves without abaxial trichomes) in wild-type, *pFD:AHL15*, *pANT:AHL15*, and *pFD:AHL15 pANT:AHL15* plants grown under SD. (**F**) and under LD (**H**). Colored dots in E-G indicate length:width ratio (E) and the number of leaves without abaxial trichomes (F,G) per plant (n = 15 biologically independent plants) per line, horizontal line and the number below this line indicate the mean and error bars indicate the s.e.m. E-G Different letters indicate statistically significant differences (P < 0.01) as determined by one-way ANOVA with Tukey’s honest significant difference post hoc test. Size bars indicate 1 cm in C and B (left panel) and 50 μm B (right panel).

Previous studies have shown that the VPC is regulated by both internal factors at the SAM (Fouracre and Scott Poethig, 2019) and peripheral organ-derived signals (Yang et al., 2011, 2013; Yu et al., 2013). During the VPC *AHL15* remains expressed in the peripheral organs, but its expression is reduced in the SAM and newly formed organs, suggesting that its expression in these tissues is most relevant for its function. To confirm this, we expressed *AHL15* under the control of the *FLOWERING LOCUS D* (*pFD*) and *AINTEGUMENTA* (*pANT*) promoters, which are predominantly active in the SAM and in young leaf primordia (Yamaguchi et al., 2016; Fouracre and Scott Poethig, 2019). Expression of *AHL15* under these promotors significantly delayed the VPC, as indicated by the reduced length/width ratio of the leaves and increased number of leaves lacking abaxial trichomes (Fig. 2C-H), and also delayed the flowering time (Supplementary Fig. 3A and B) under both SD and LD conditions. Combining both *pFD:AHL15* and *pANT:AHL15* constructs in one plant line led to a further delay of the VPC (Fig. 2C-H) and flowering time (Supplementary Fig. 3A and B). This additive effect can be explained by the slightly different but overlapping activities of the selected promoters (Yamaguchi et al., 2016; Fouracre and Scott Poethig, 2019). These results indicate that *AHL15*-induced delay in the VPC and flowering time is achieved through its expression in the SAM and leaf primordia, and not by its expression in older leaves (Fig. 2B).

### AHL proteins repress *SPL* gene expression in a miR156/miR157-independent manner

In Arabidopsis, the VPC is mediated by a gradual decrease in *miR156*/*miR157* expression, which increases the expression of the *SPL* miR156/miR157 target genes. *SPL* genes in turn promote the adult developmental program in the SAM, resulting in the production of adult instead of juvenile leaves and eventually in the initiation of flowering (Wang et al., 2009; Wu et al., 2009). One possible mode of action of AHL proteins might be that they suppress *SPL* expression by enhancing the *miR156*/*miR157* pathway. A comparison of the expression of several *SPL* genes (*SPL2, SPL9*, *SPL10*, *SPL11*, *SPL13*, and *SPL15*) known to promote the VPC (Xu et al., 2016) by qPCR showed that the transcript levels of *SPL2*, *SPL9*, *SPL13*, and *SPL15* were significantly upregulated in shoots of 3-week-old *ahl15/+ pAHL15:AHL15-ΔG* mutant compared to wild-type seedlings (Fig. 3A), and that only *SPL2* expression was higher 2-week-old mutant seedling shoots (Fig. 3A). This is in line with the timing of the VPC and the dynamics of *AHL* gene expression (Fig 2A, B), and indicates that AHL proteins repress the expression of *SPL2*, *SPL9*, *SPL13*, and *SPL15* during this period.

**Figure 3.**
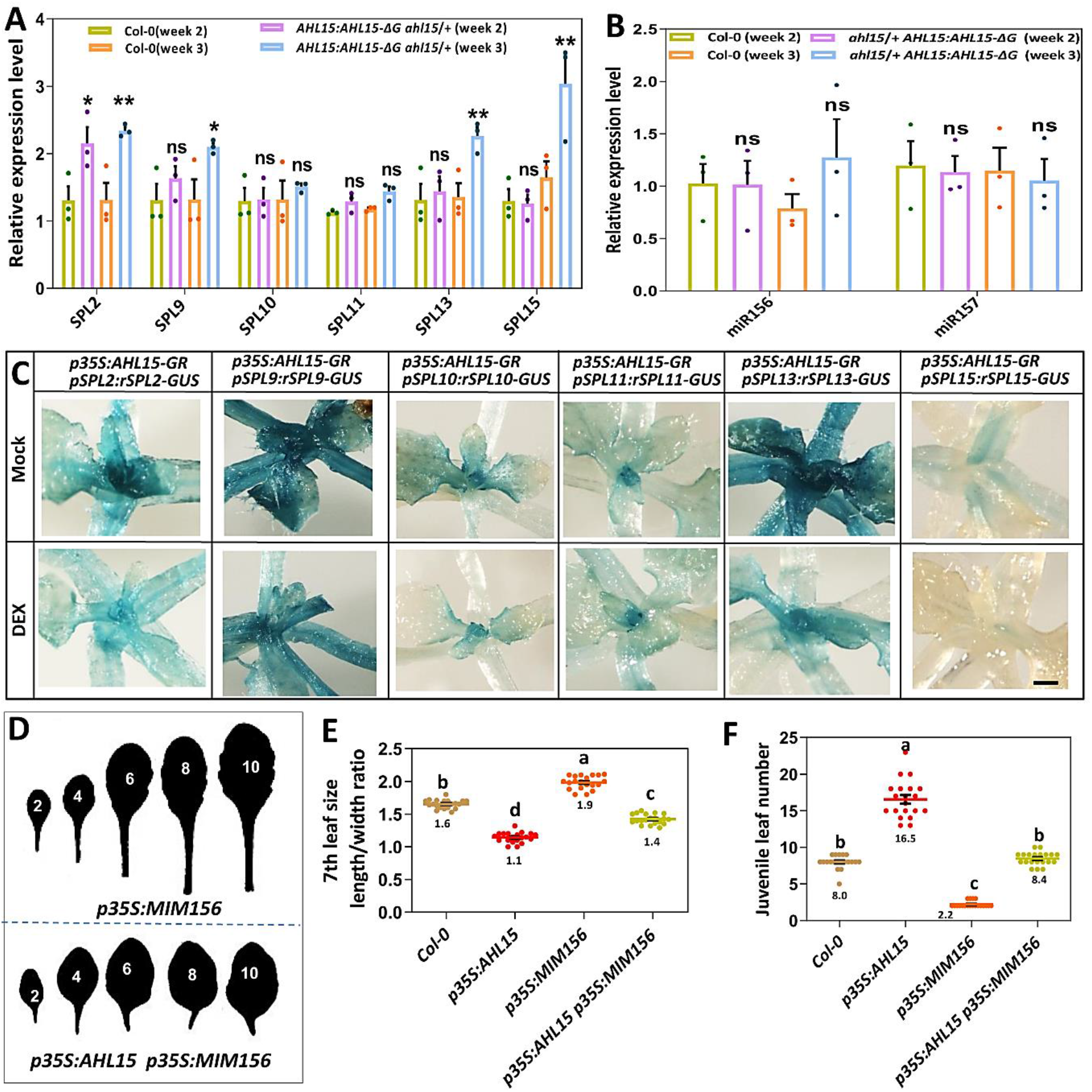
AHL15 represses *SPL* gene expression in a miRNA-independent manner. (**A, B**) The relative expression level of *SPL2, SPL9*, *SPL10*, *SPL11*, *SPL13*, and *SPL15* (**A**) or *miR156* and *miR157* (**B**) in the shoot apex and young leaves of 2- and 3-week-old wild-type and *ahl15/+ pAHL15:AHL15-ΔG* plants grown under SD conditions. (Dots indicate the values of three biological replicates per plant line, the bar indicates the mean and error bars indicate the s.e.m. Asterisks indicate significant differences from wild type (Col-0, *P < 0.05, **P < 0.01, ns = not significant or P ≥ 0.05), as determined by a two-sided Student’s *t*-test. In **B,** values were normalized to the value in wild-type in week 2. (**C**) Histochemical staining for GUS activity in 10-day-old shoot apex and young leaves of water (mock) or 20 μM DEX (+DEX) treated (for 2 days) *p35S:AHL15-GR* plants expressing the miR156 resistant *pSPL9:rSPL9-GR*, *pSPL2:rSPL2-GUS*, *pSPL9:rSPL9-GUS, pSPL10:rSPL10-GUS, pSPL11:rSPL11-GUS, pSPL13:rSPL13-GUS* or *pSPL15:rSPL15-GUS* reporter grown under LD conditions. Size bars indicate 1 mm. (**D**) Shape of the successive rosette leaves of 7-week-old *p35S:MIM156* or *p35S:AHL15 p35S:MIM156* plants grown under SD conditions. (**E, F**) The length:width ratio of the 7 th leaf (**E**) or The juvenile leaf number (leaves without abaxial trichomes) (**F**) of 7-week-old wild-type, *p35S:MIM156*, *p35S:AHL15* or *p35S:AHL15 p35S:MIM156* plants grown under SD conditions. Colored dots indicate length:width ratio (E) or the number of leaves without abaxial trichomes (F) per plant (n = 15 biologically independent plants) per line, horizontal line and the number below this line indicate the mean and error bars indicate the s.e.m. Different letters indicate statistically significant differences (P < 0.01) as determined by a one-way ANOVA with Tukey’s honest significant difference post hoc test.

The fact that not all six *SPL* genes tested were simultaneously upregulated in the *ahl15/+ pAHL15:AHL15-ΔG* mutant background suggested that AHLs repress *SPL* expression independent of the *miR156*/*miR157* pathway. Indeed, *pmiR156A:GUS*, *pmiR156B:GUS*, and *pmiR156D:GUS* reporters (Yu et al., 2015) did not show a major change in expression upon AHL15 activation by DEX treatment in the *p35S:AHL15-GR* background (Supplementary. Fig, 4A). Also qPCR analysis showed that miR156/157 levels were not significantly different in 2-or 3-week-old wild-type or *ahl15/+ pAHL15:AHL15-ΔG* seedlings grown under juvenile phase prolonging SD conditions (Fig. 3B). In addition, when testing the expression of six miR156/miR157-insensitive *pSPL:rSPL-GUS* reporters (e.g., *rSPL2*, *rSPL9*, *rSPL10*, *rSPL11*, *rSPL13*, *rSPL15*) (Xu et al., 2016) in the *p35S:AHL15-GR* background, the expression of *rSPL2*, *rSPL9*, *rSPL13*, and *rSPL15* appeared to be down-regulated by DEX treatment (Fig. 3C). Together these results indicate that AHL proteins suppress the expression of specific *SPL* genes involved in the VPC in a miR156/miR157-independent manner.

In plants overexpressing a target mimic of *miR156* (*p35S:MIM156*) *SPL* expression is enhanced (Franco-Zorrilla et al., 2007), resulting in the accelerated appearance of adult leaves (Fig. 3D-F) and early flowering (Supplementary Fig. 5A and B). *AHL15* overexpression negated the precocious appearance of adult vegetative traits (Fig. 3D-F) and early flowering (Supplementary Fig. 5A and B) of *p35S:MIM156* plants, bringing these traits back to or close to wild-type levels. In the reverse experiment where the *p35S:AHL15* construct was introduced into the *p35S:miR156* background having low SPL levels, we observed a remarkably additive effect on the vegetative phase transition and flowering time (Supplementary Fig. 6A and B). The results of both experiments support our model that AHL15 and family members suppresses *SPL* gene expression in a miR156/miR157-independent manner.

### SPLs promote the vegetative phase change in part by repressing *AHL15* expression

During the VPC in 2-week-old Arabidopsis seedlings, increasing SPL levels (Wang et al., 2009) coincided with downregulation of *AHL15*, *AHL19* and *AHL20* (Fig 1A, B). This led us to hypothesize that the SPL transcription factors are mediating down-regulation of *AHL* gene expression. Based on this hypothesis, we expected *AHL* expression to be downregulated in Arabidopsis lines with elevated SPL levels (e.g. *p35S:MIM156* or *pSPL9:rSPL9*) and to be upregulated in Arabidopsis lines with reduced SPL levels (e.g. *p35S:miR156* or *spl* loss-of-function mutants). qPCR analysis indeed showed that the expression of *AHL15* and *AHL20* was significantly reduced in one-week-old *p35S:MIM156* seedlings compared to wild-type seedlings (Fig. 4A). In line with this result, expression of the *pAHL15*:*GUS* reporter was reduced in one-week-old seedlings of the *p35S:MIM156* line compared to wild-type seedlings (Fig. 4B), or in DEX-treated *pSPL9:rSPL9-GR* seedlings compared to mock-treated seedlings (Fig. 4C). In contrast, expression of the *pAHL15:GUS* reporter seemed rather enhanced in two-week-old *p35S:miR156* seedlings compared to wild-type seedlings (Fig. 4D).

**Figure 4.**
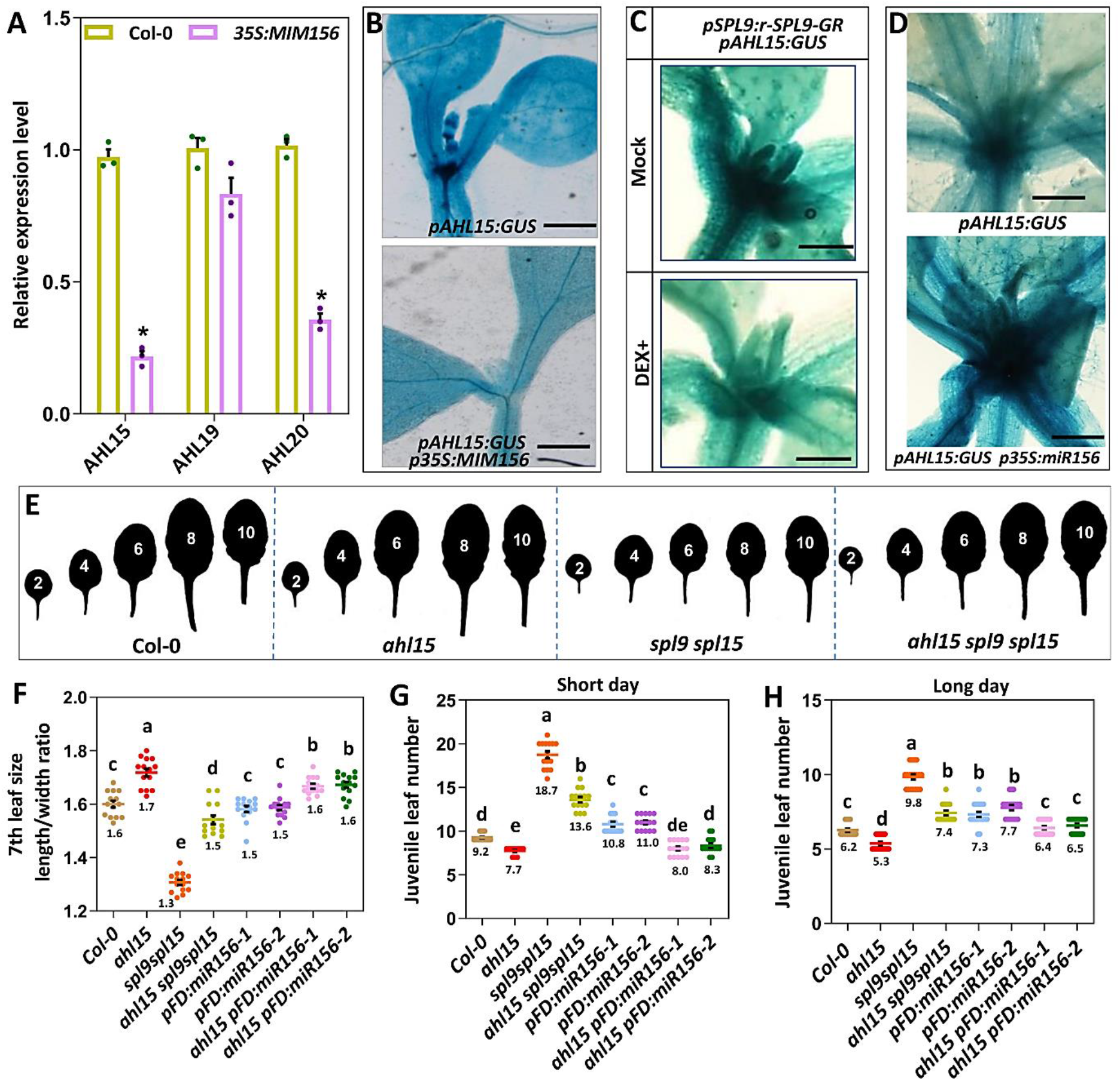
SPLs promote the VPC by repressing *AHL15* expression. (**A**) Relative expression level of *AHL15*, *AHL19* and *AHL20* in the shoot apex and young leaves of 2-week-old wild-type or *p35S:MIM156* seedlings grown under LD conditions. Dots indicate the values of three biological replicates per plant line, the bar indicates the mean, and error bars the s.e.m Asterisks indicate a significant difference (p < 0.01), as determined by a two-sided Student’s *t*-test. (**B-D**) Histochemical staining for GUS activity in 8-day-old *pAHL15:GUS* (top) or *p35S:MIM156 pAHL15:GUS* (bottom) seedlings (**B**), 10-day-old water (mock) or 20 μM DEX (+DEX) treated *pAHL15:GUS p35S:SPL9-GR* seedlings (**C**), or 15-day-old wild-type (top) or *p35S:miR156* (bottom) seedlings (**D**) grown under LD conditions. (**E**) Shape of the successive rosette leaves of 7-week-old wild-type, *ahl15*, *spl9 spl15* and *ahl15 spl9* spl15 plants grown under SD conditions. Size bars indicate 1 mm. (**F**) The length:width ratio of the 7th leaf of 7-week-old wild-type, *ahl15*, *spl9 spl15* and *ahl15 spl9* spl15 plants grown under SD conditions. (**G**, **H**) The juvenile leaf number (leaves without abaxial trichomes) in wild-type, *ahl15*, *spl9 spl15* and *ahl15 spl9* spl15 plants grown under SD (**G**) or LD (**H**) conditions. Colored dots in **F** - **H** indicate length:width ratio of the 7th leaf (**F**) or the number of leaves without abaxial trichomes (**G**,**H**) per plant (n = 15 biologically independent plants) per line, horizontal line and the number below this line indicate the mean and error bars indicate the s.e.m. Different letters indicate statistically significant differences (P < 0.01) as determined by one-way ANOVA with Tukey’s honest significant difference post hoc test.

Based on our hypothesis, the delayed VPC and flowering phenotypes of plants with reduced SPL levels (e.g. the *spl9 spl15* double loss-of-function mutant and *p35S:miR156*; Wu et al., 2009) would be dependent on the elevated expression of functional *AHL* genes. The *ahl15* loss-of-function mutation in the *ahl15 spl9 spl15* triple mutant indeed partially rescued the delayed VPC phenotypes of the *spl9 spl15* double mutant. The length/width ratio of *ahl15 spl9 spl15* leaves was almost restored to wild-type levels, and the VPC of *ahl15 spl9 spl15* plants was significantly accelerated compared to that of *spl9 spl15* double mutant plants, both in SD and in LD conditions (Fig. 4E-G). Introduction of the *p35S:miR156* line in the *ahl15* loss-of-function background also rescued the delay in VPC induced by *miR156* overexpression back to wild type levels, even in heterozygous *ahl15/+ 35S:miR156* plants. Since we could not exclude that the T-DNA insertion in the *ahl15* mutant silenced the *p35S:miR156* construct (Daxinger et al., 2008), we placed the *miR156* gene under control of the *FD* promotor (*pFD:miR156*) and introduced the resulting construct into the *ahl15* mutant and wild-type background. The delay in VPC caused by *FD* promotor-controlled *miR156* expression in the wild-type background was significantly reduced in the *ahl15* background both under SD and LD conditions (Fig. 4F-H). These results indicate that a functional *AHL15* gene is required for the delay in VPC in plants with reduced SPL levels. Together the data confirms our hypothesis that the repression of *AHL*s in the SAM and flower primordia is mediated by SPLs, either indirectly or directly by binding of these transcription factors to the *AHL* regulatory regions.

In contrast to the VPC results presented above, the *ahl15* loss-of-function mutation had no or only a marginal effect on the delayed flowering of the *spl9 spl15* or *pFD:miR156* plants (Supplementary Fig. 7). It should be noted, however, that *ahl15* single or *ahl15 ahl19* double mutant plants show a wild-type flowering time, and that only *ahl15 ahl19 p35S:amiRAHL20* triple mutant or the *ahl15/+ pAHL15:AHL15-ΔG* dominant negative mutant plants flower significantly earlier compared to wild type (Supplementary Fig. 1, Karami et al., 2020a).This indicates that not *AHL15* but other redundantly acting *ALH* genes, such as *AHL20* (Fig 4A), are the prime targets for repression by SPLs in the promotion of flowering.

### SPLs promote reproductive identity of axillary meristems by repressing *AHL15* expression

Our previous results have shown that *AHL15* does play a central role in axillary meristem (AM) maturation, i.e. the switch from vegetative to reproductive identity (Karami et al., 2020a). Overexpression of *AHL15* (*p35S:AHL15* or *pMYB:AHL15*) repressed AM maturation, leading to their prolonged vegetative activity and resulting in the formation of aerial rosettes from inflorescence nodes. Such aerial rosettes are not formed on wild-type Arabidopsis (Col-0) plants grown under LD conditions (Supplementary Fig. 8A and B), but can be induced by growing plants under SD conditions or by mutating the *SOC1* and *FUL* genes encoding transcription factors that suppress *AHL15* expression. We have demonstrated that the aerial rosette phenotype is dependent on a functional *AHL15* gene (Karami et al., 2020a). Interestingly, the AMs in the axils of cauline leaves of the *spl9 spl15* mutant or *p35S:miR156* (or *pFD:miR156*) overexpression plants also produced aerial rosettes (Supplementary Fig. 8A and B), revealing a yet unidentified role for SPLs in promoting the vegetative to reproductive transition of AMs. Introduction of *spl9 spl15* or *pFD:miR156* in the *ahl15* loss-of-function mutant background strongly reduced the aerial rosette formation phenotype of both mutant lines (Fig. 5A and B), indicating that *AHL15* function is important for the frequent aerial rosette formation in *spl9 spl15* and *pFD:miR156* plants, which is completely absent in wild-type and *ahl15* plants when grown under the same LD conditions (Karami et al., 2020a). In line with the role of the SPL transcription factors as *AHL* repressors, *AHL15* expression was significantly increased in *spl9 spl15* and *p35S:miR156* AMs compared to wild-type AMs (Fig. 5C and D). Based on these results, we concluded that *AHL15* acts downstream of the *SPL* genes, and that repression of *AHL15* by SPLs in wild-type plants promotes the reproductive identity of AMs, whereas under conditions where SPL activity is reduced (e.g. in *p35S:miR156* or *spl9 spl15* plants) elevated *AHL15* expression enhances the vegetative activity of AMs, resulting in the formation of aerial rosettes.

**Figure 5.**
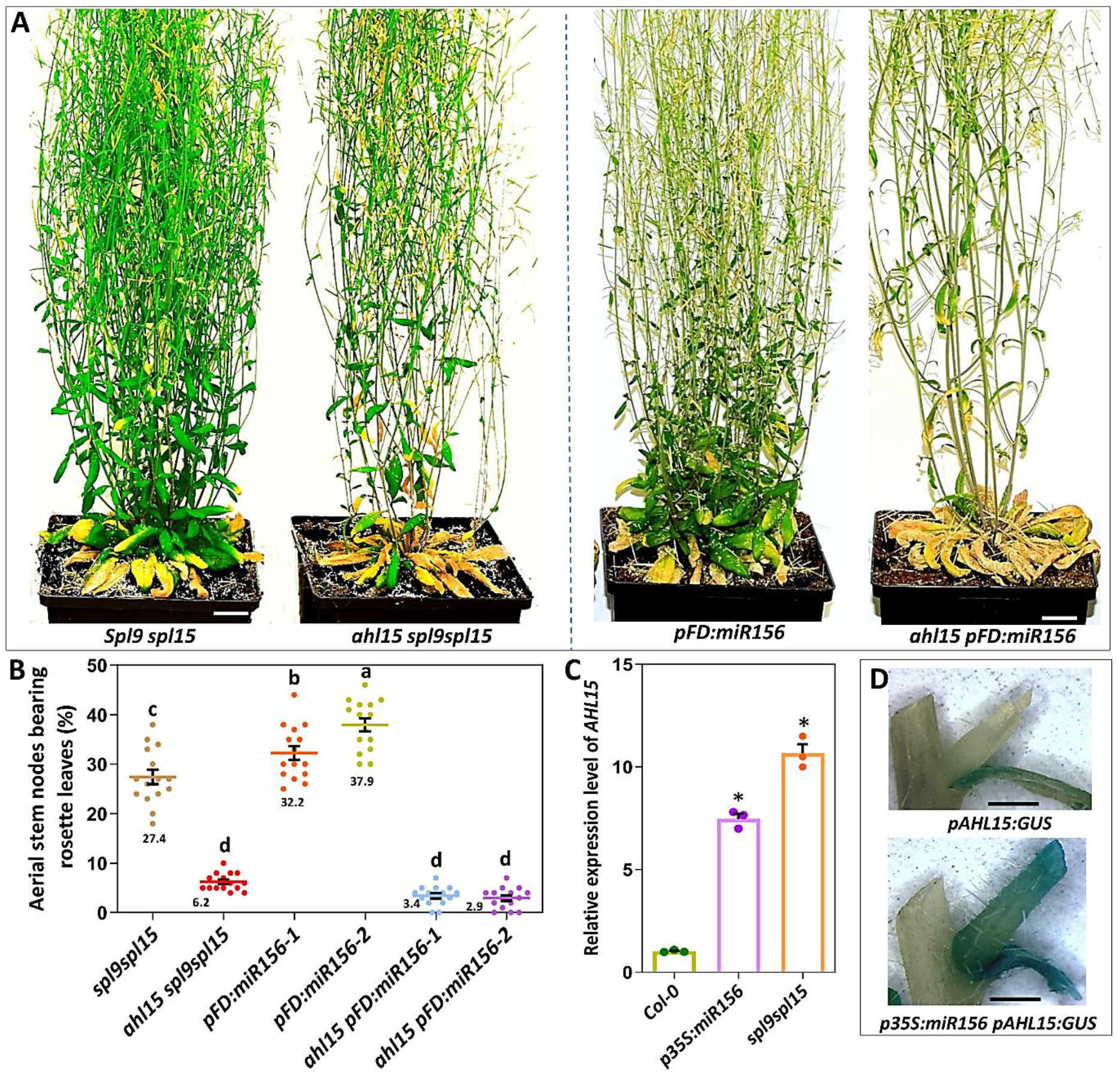
SPLs promote reproductive identity of axillary meristems by repressing *AHL15*. (**A**) Phenotype of 3-month-old *spl9 spl15*, *ahl15 spl9 spl15*, *pFD:miR156* or *pFD:miR156 ahl15* plants grown under LD conditions. (**B**) Percentage of the aerial stem nodes bearing rosette leaves in 3-month-old \*spl9 spl15*, *ahl15 spl9 spl15*, *pFD:miR156*, *ahl15 pFD:miR156* plants grown under LD conditions. Please note that wild-type and *ahl15* stem nodes do not form rosette leaves under these conditions. Colored dots indicate the percentage per plant (n = 15 biologically independent plants) per line, horizontal line indicates the mean and error bars indicate the s.e.m. Different letters indicate statistically significant differences (P < 0.01) as determined by one-way ANOVA with Tukey’s honest significant difference post hoc test. (**C**) The relative expression level of *AHL15* in aerial stem nodes of wild-type, *p35S:miR156* or *spl9 spl15* plants 2 weeks after flowering. Dots indicate the values of three biological replicates per plant line, horizontal line and the number below this line indicate the mean and error bars indicate the s.e.m. The asterisk indicates a significant difference (p < 0.01), as determined by a two-sided Student’s *t*-test. (**C**) Histochemical staining for GUS activity in inflorescence nodes of six-week-old wild-type and *p35S:miR156* plants grown under LD conditions. Size bars indicate 1 cm in A and 50 mm in D.

## Discussion

Plants during their lifetime progress through distinct consecutive developmental phases, starting with embryogenesis and followed by respectively the juvenile-, adult- and reproductive phase. What drives and regulates the transition from one developmental phase to another is a longstanding fundamental question in plant developmental biology. In Arabidopsis and several other plants, the transition from the juvenile to adult phase (VPC) and from the adult to reproductive phase (flowering) has been shown to be driven by the ageing pathway. In this pathway, the SPL transcription factors promote phase transitions, and miR156 and miR157 repress these transitions by targeting the *SPL* transcripts (Poethig, 2013; Teotia and Tang, 2015). Previously we have shown that Arabidopsis *AHL15* and its paralogs enhance plant longevity by delaying maturation of AMs (Karami et al., 2020a). Here we show that AHL proteins interfere with the ageing pathway by repressing *SPL* gene expression in a miRNA-independent manner, and that in turn *AHL15* expression is repressed through feedback regulation by the SPLs. In addition, we show that this reciprocal negative feedback loop between *AHL* and *SPL* genes not only regulates the VPC and flowering but also AM maturation and thus controls plant longevity and -life history.

### *AHL* genes delay the VPC

Although the adult vegetative to reproductive phase change is agronomically most important, the VPC also has a strong impact on plant fitness and biomass, not in the least because it determines the balance between vegetative growth and reproductive development (Demura and Ye, 2010). Although the timing of the VPC can be influenced by environmental factors such as photoperiod, light intensity and temperature, this developmental switch is mainly regulated by endogenous genetic components (Poethig, 2013). Studies in Arabidopsis have revealed that the gradual decline in the expression of miR156/miR157 increases the abundance of the SPL transcription factors, which promote the VPC (Poethig, 2013; Teotia and Tang, 2015). The change in leaf morphology during the VPC is accompanied by a reduced expression of *AHL15* and close homologs in the SAM and young leaves, which is in line with the repression of *AHL* genes by SPLs.

Recently, it has been reported that these internal factors at the SAM maintain the juvenile phase during early shoot development, but that the SAM plays a relatively minor role in the regulation of leaf identity at later stages of the VPC (Fouracre and Poethig, 2019). The fact that *ahl* loss-of-function plants (*ahl15/+ pAHL15:AHL15-ΔG*) immediately form adult leaves, and that *p35S:AHlL15* seedlings form multiple juvenile leaves confirm that AHLs belong to the internal factors in the SAM that maintain the juvenile phase in early shoot development. However, when we overexpressed *AHL15* specifically in young leaf primordia this also delayed the VPC, which is in line with the observation that the VPC is regulated by peripheral organ-derived signaling (Yang et al., 2011, 2013; Yu et al., 2013) and that AHL15 can control the VPC both internally at the SAM and through leaf-derived signalling (Poethig, 2013; Teotia and Tang, 2015).

Our analysis showed that AHL15 represses the expression of *SPL* genes in a miR156/miR157-independent manner. Previous studies have shown that the abundance of miR156/miR157 is highly declined (about 90%) at the shoot apex of Arabidopsis seedlings in a two weeks period, whereas the level of the most SPLs remained constant or even slightly increased during this period (Xu et al., 2016). We showed that the level of *SPL2, SPL9*, *SPL13*, and *SPL15* are significantly higher at the shoot apex of *ahl15* loss-of-function compared to wild-type plants, whereas the level of miR156/miR157 did not change between wild-type or *ahl15* loss-of-function mutant plants. This clearly indicates that the maintained repression of SPL expression is mediated by AHL15 independent of miR156/157. How AHL15 represses *SPL* genes remains unknown. AHL15 might directly bind to the *SPL* loci. However, preliminary yeast one-hybrid assays did not provide an indication for direct binding of AHL15 to the promoter regions of *SPL* (data not shown). Recently, we have shown that AHL15 overexpression suppresses the biosynthesis of the plant hormone gibberellic acid (GA) (Karami et al., 2020a). Since DELLA proteins, the degradation targets of GA signalling, have been shown to repress *SPL* expression (Yu et al., 2012), AHL15 may repress *SPLs* by reducing GA biosynthesis and thus stabilizing the DELLA proteins (Supplementary Figure 9).

Previously, it has been shown that SPLs promote trichome development on the abaxial side of leaves partially by repressing the AP2-like transcription factors TOE1 and TOE2 (Wu et al., 2009). How SPLs promote the other adult leaf traits such as leaf elongation and leaf serration was not known until now. In this study, we showed that the promotion of adult traits, including leaf elongation and trichomes on the abaxial side of leaves, by SPLs is contributed by the repression of *AHL* genes, as the delay in the juvenile-to-adult transition by miR156 requires AHL15. Thus, we suggest a reciprocal negative feedback loop between SPL and AHL (Fig. 6), and although we do not know whether AHLs and SPLs regulate each other’s expression directly or indirectly, our findings provide a new advance in the understanding of the regulation of VPC.

**Figure. 6.**
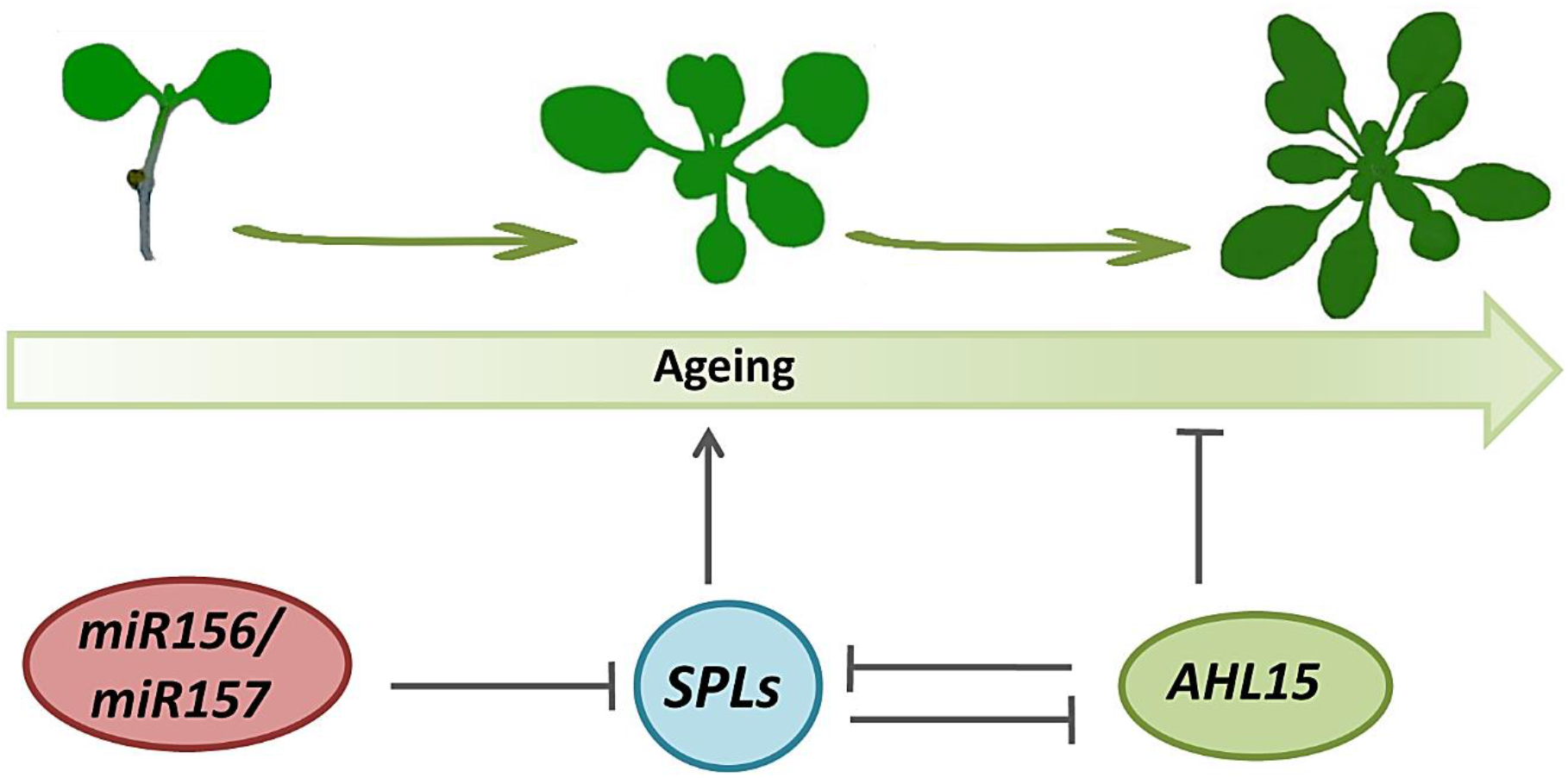
Proposed model for the role of AHL15 and family members in the regulation of plant ageing. Following their germination, high expression of *miR156* and *AHL15* and family members keeps seedlings in the juvenile vegetative phase by suppressing the expression of the ageing promoting SPL transcription factors. When seedlings get older, gradual down-regulation of *miR156* elevates SPL abundance, which in turn promotes the juvenile to adult phase transition (VPC) by stimulating plant ageing in part by downregulating AHLs. Negative feed-back between AHLs and SPLs moderates plant ageing. Arrows indicate activation, and blunted lines indicate repression.

### *AHL* genes delay flowering time

The switch from vegetative growth to flowering is a major developmental transition in plants. Two decades of genetic and physiological studies have led to the identification a number of pathways controlling flowering, including the photoperiod pathway, the vernalization pathway, the GA pathway, and the age pathway (Supplementary Fig. 9) (Turnbull, 2011; Matsoukas et al., 2012; Song et al., 2013). In this work, we have revealed that *AHL15* and close homologs are involved in the control of flowering time. We showed that overexpression *AHL15* causes extremely late flowering. The late-flowering phenotypes were also observed in overexpression of *AHL18*, *AHL22*, *AHL27*, and *AHL29* (Street et al., 2008; Xiao et al., 2009; Yun et al., 2012; Xu et al., 2013). However, these overexpression phenotypes did not reflect the normal function of these genes since no effect or relatively small effects were on flowering time observed in single loss-of-function mutants of these genes. Thus, the role of these AHLs and other members of the AHL family in the control flowering time has remained elusive until now. Our results showed, however, that loss-of-function of *AHL15* and close homologs does promote flowering time, indicating an important role for these genes in regulating the vegetative to reproductive phase change.

An increase in SPL proteins levels promotes the flowering time by activating the transcription of *SOC1* and several other floral-promoting genes at the SAM (Wang et al., 2009; Yu et al., 2012). Consistent with delay in the flowering time by targeted expression of *AHL15* in the SAM, we postulate that AHL15-suppressed flowering time at the SAM is contributed by the repression of SPLs at the SAM (Supplementary Fig. 9). FLOWERING LOCUS T (FT) that is produced in leaves and is transported as a florigen signal from the leaves to the SAM (Supplementary Fig. 9), triggers the flowering by activating the transcription of several flowering-promoting genes (Abe et al., 2005; Corbesier et al., 2007). It has been shown that overexpression of *AHL22* delays flowering time in Arabidopsis by repressing *FT* in leaves (Yun et al., 2012). Consistent with delay in the flowering time by targeted expression of AHL15 in leaves, the AHL15-suppressed flowering time through leaves is might be also contributed to repression *FT* (Supplementary Fig. 9). Interestingly, the MADS box transcription factor *FLOWERING LOCUS C* (*FLC*) which acts as a potent repressor of flowering time could also suppress the *FT* and *SPL* genes (Deng et al., 2011; Matsoukas et al., 2012). Therefore, a molecular link between AHL and FLC in the repression of the *SPLs* and *FT* is predicted (Supplementary Fig. 9). GA is known to accelerate flowering by degradation of DELLA proteins (Yu et al., 2012). We showed that AHL15 reduces transcription of genes that encode the rate-limiting enzyme in the synthesis of GA biosynthesis (Karami et al., 2020a), therefore, regulation of GA biosynthesis by AHL15 is another possibility of suppressing the flowering time by AHL15 (Supplementary Fig. 9). Taken together, we suggest that AHL proteins act as flowering repressors by down-regulating many genes in different pathways controlling flowering (Supplementary Fig. 9). More detailed investigation of the relationship between AHL proteins and the different flowering pathways is required to elucidate which interactions are truly relevant.

### SPLs promote reproductive identity of axillary meristems by repressing *AHL15* expression

In Arabidopsis, following the floral transition the main inflorescence meristem (IM) is converted into a floral meristem after producing two to three cauline leaves. The AMs formed in axils of these cauline leaves directly become IM that produce a few lateral cauline leaves before being converted to a floral meristem (Supplementary Fig. 8). In contrast, the AMs in the axils of cauline leaves of *p35S:miR156* or *spl9 spl15* and *p35S:AHL15* plants start in the vegetative phase and are converted to IM only after producing several rosette leaves (Supplementary Fig. 8). This heterochronic shift in the development of AMs leading to the formation of aerial rosettes in *p35S:miR156* or *spl9 spl15* plants is associated with the up-regulation of *AHL15*. Moreover, we have previously shown that the development of aerial rosettes from AMs in axils of cauline leaves of the *soc1 ful* double mutant (Melzer et al., 2008) is associated with and dependent on upregulation of *AHL15* expression (Karami et al., 2020a). Consistent with the finding that SPL proteins promote *SOC1* and *FUL* expression (Wang et al., 2009) and our own results, we postulate that SPLs prevent the aerial rosettes formation by upregulating *SOC1* and *FUL*, which subsequently leads to repression of *AHL15* expression (Supplementary Fig. 9).

A consequence of maintaining AMs in the vegetative phase, following the floral transition in *p35S:miR156*, *spl9 spl15*, *p35S:AHL15* or *soc1 ful* plants, is that this increases the life span of the plant and extends the period during which a plant is able to produce fruits and seeds. This heterochronic growth and development of AMs has been reported in Arabidopsis ecotypes such as Sy-0 (Poduska et al., 2003). A dominant allele of *FLC* has been reported as a key factor that underlies the aerial-rosette phenotype and increases the life span in Sy-0 (Poduska et al., 2003). Detailed studies on the genetic link between *FLC* and *AHL*s should provide more insight into aerial-rosette phenotype in Arabidopsis ecotypes such as Sy-0. *AHL* gene families have been identified in many plant species (Zhao et al., 2014; Karami et al., 2020b). Along with the results obtained in tobacco, we suggest that the role of *AHL* genes in controlling developmental switches and through that plant longevity and life history strategy is likely to be conserved to in many other plants.

## Methods

### Plant material and growth conditions and phenotype analysis

All Arabidopsis mutant- and transgenic lines used in this study are in the Columbia (Col-0) background. The *ahl15*, *ahl19*, *p35S:AHL15*, *p35S:AHL15-GR*, *pAHL15:GUS*, *pAHL15:AHL15*, *pAHL15:AHL15-ΔG*, *p35S:amiRAHL20 and ahl15/+ pAHL15:AHL15-ΔG* plant lines has been described previously (Karami et al., 2020a,b). The *spl9 spl15, p35S:miR156*, *p35S:MIM156*, *pSPL9:rSPL9-GR*, *pSPL2:rSPL2-GUS*, *pSPL9:rSPL9-GUS, pSPL10:rSPL10-GUS, pSPL11:rSPL11-GUS, pSPL13:rSPL13-GUS* and *pSPL15:rSPL15-GUS* plant lines were obtained from the Nottingham Arabidopsis Stock Centre (NASC). The reporter lines *pmiR156A:GUS, pmiR156B:GUS and pmiR156C:GUS* has been described previously (Yu et al., 2015). Plant lines and F1, F2 or F3 plants from crosses were PCR genotyped using primers described in the Supplementary Table 1. Seeds were directly sown on soil in pots and grown at 21°C, 65% relative humidity, and 16 hours (long day: LD) or 8 hours (short day: SD) photoperiod. One exception: SD was10 hours for the top panel of supplementary Fig. 1A. To score aerial rosette leaves production by AMs, wild-type, mutant or transgenic plants were transferred to larger pots about 2 weeks after flowering. *Nicotiana tabacum* cv SR1 Petit Havana (tobacco) wild-type or *p35S:AHL15-GR* plants (Karami et al., 2020a) were grown in medium-sized pots at 25 °C, 70% relative humidity, and a 16 hours photoperiod. For dexamethasone (DEX, Sigma-Aldrich) treatment, Arabidopsis and tobacco plants were sprayed with 20 and 30 μM DEX, respectively. Leaf size was measured directly using a ruler.

The number of juvenile leaves (without abaxial trichomes) was scored once they appeared, and detected by eye. For imaging of leaf shape, fully expanded leaves were removed, attached to cardboard with double-sided tape, flattened and photographed with a Nikon D5300 camera. Leaf images were optimized and changed into black and white images and assembled using Adobe Illustrator cc2017. Potted plants were photographed with a Nikon D5300 camera. All measurements were statistically analyzed and plotted into graphs in GraphPad Prism 8.

### Plasmid construction and transgenic Arabidopsis lines

To generate the constructs *pFD:AHL15* and *pANT:AHL15*, 3 kb regions upstream of the ATG initiation codon of the *FD* (AT4G35900) and *ANT* (AT4G37750) genes were amplified from ecotype Columbia (Col-0) genomic DNA using the forward (F) and reverse (R) PCR primers indicated in Supplementary Table S1. The resulting fragments were first inserted into pDONR207 by BP reaction, and subsequently cloned upstream of the genomic fragment containing the *AHL15* transcribed region in destination vector *pGW-AHL15* (Karami et al., 2020b) by LR reaction. To generate the *pFD:miR156*, first *pGW-miR156* was generated by replacing the *AHL15* fragment in *pGW-AHL15* for a *Kpn*I and *Spe*I fragment containing the *miR156* transcribed region. The *FD* promoter fragment was subsequently recombined from pDONR207 into the resulting *pGW-miR156* construct by LR reaction. All binary vectors were introduced into *Agrobacterium tumefaciens* strain AGL1 by electroporation (den Dulk-Ras and Hooykaas, 1995) and Arabidopsis Col-0 and *ahl15* plants were transformed using the floral dip method (Clough and Bent, 1998). The resulting plant lines and F1, F2 or F3 plants from crosses were PCR genotyped using primers described in the Supplementary Table 1.

### Histochemical staining and microscopy

Histochemical staining of plant tissues for β-glucuronidase (GUS) activity was performed as described previously (Anandalakshmi et al., 1998). Tissues were stained for 4 hours at 37°C, followed by chlorophyll extraction and rehydration by incubation for 10 minutes in a graded ethanol series (75, 50, and 25 %). GUS stained tissues were observed and photographed using a LEICA MZ12 microscope (Switzerland) equipped with a LEICA DC500 camera.

### Quantitative real-time PCR (qPCR) analysis

RNA was isolated from rosette base nodes, and the basal part of inflorescence stems (about 0.5 cm above rosette base) using the RNEasy© kit (Qiagen). First-strand cDNA was synthesized using the RevertAid RT Reverse Transcription kit (Thermo Fischer Scientific). Quantitative PCR was performed on three biological replicates along with three technical replicates using the SYBR-green dye premixed master-mix (Thermo Fischer Scientific) in a C1000 Touch© thermal cycler (BIO-RAD). CT values were obtained using Bio-Rad CFX manager 3.1. The relative expression level of genes was calculated according to the 2-ΔΔCt method (Livak and Schmittgen, 2001).

The miRNAs abundance was quantified using 1μg of RNA in a reverse transcription reaction by using SnoR101 reverse primer and a miRNA-specific RT primer. Expression was normalized using the *β-TUBULIN-6* gene, analyzed and plotted into graphs in GraphPad Prism 8. Three biological replicates were performed, with three technical replicates each. The primers used are described in Supplementary Table 1.

## Acknowledgements

We are grateful to Scott Poethig for providing different *SPL* and *miR156* related lines also Nam-Hai Chua for providing *pmiR156-GUS* reporter lines. We are grateful to Ward de winter, Jan Vink, Nick Surtel and Mariel Lavrijsen for their technical supports.

## Author contribution

A.R., O.K. and R.O. conceived the project, designed the experiments and analysed and interpreted the results. A.R. performed all the experiments. O.K. and R.O. supervised the project. A.R., O.K. and R.O. wrote the manuscript. All authors read and commented on versions of the manuscript.

---

**Supplementary Fig. 1.**
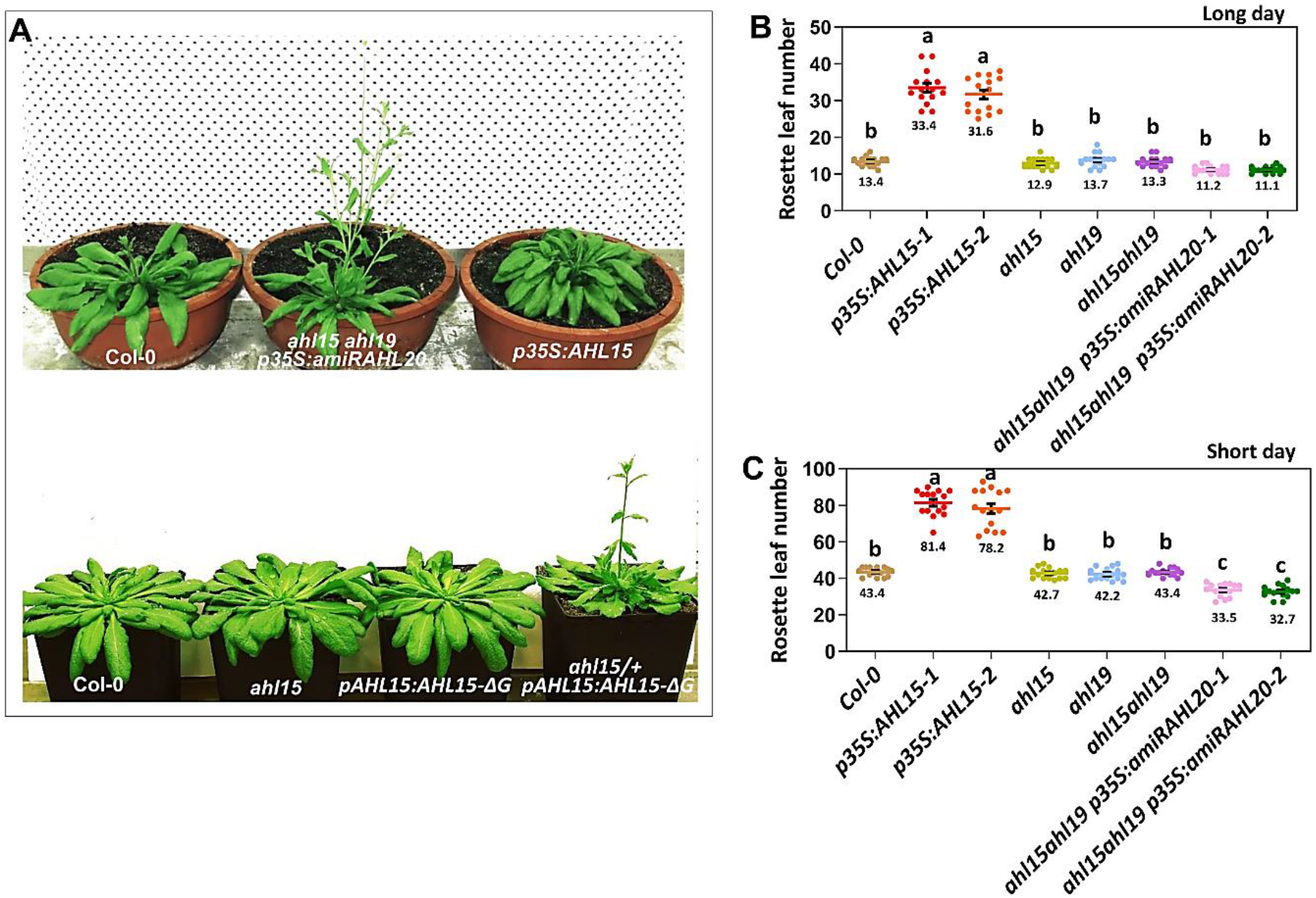
*Arabidopsis AHL15* and close homologs redundantly regulate flowering time. (**A**) Phenotype of a 8-weeks-old wild-type (Col-0), *ahl15 ahl19 p35S:amiRAHL20* or *p35S:AHL15* plant grown under SD (10-hour photoperiod) conditions (top) or of a 10-weeks-old wild-type (Col-0), *alh15*, *pAHL15:AHL15-ΔG* or *ahl15/+ pAHL15:AHL15-ΔG* plant grown under SD (8-hour photoperiod) conditions (bottom). (**B**, **C**) The number of rosette leaves produced until flowering by wild-type (Col-0), *p35S:AHL15, ahl15*, *ahl19, ahl15 ahl19*, or *ahl15 ahl19 p35S:amiRAHL20* plants grown under LD (**B**) or SD (**C**) conditions. Colored dots indicate the number of rosette leaves per plant (n = 15 biologically independent plants) per line, horizontal line and the number below this line indicate the mean and error bars indicate the s.e.m. Different letters indicate statistically significant differences (P < 0.01) as determined by a one-way ANOVA with Tukey’s honest significant difference post hoc test.

**Supplementary Fig. 2.**
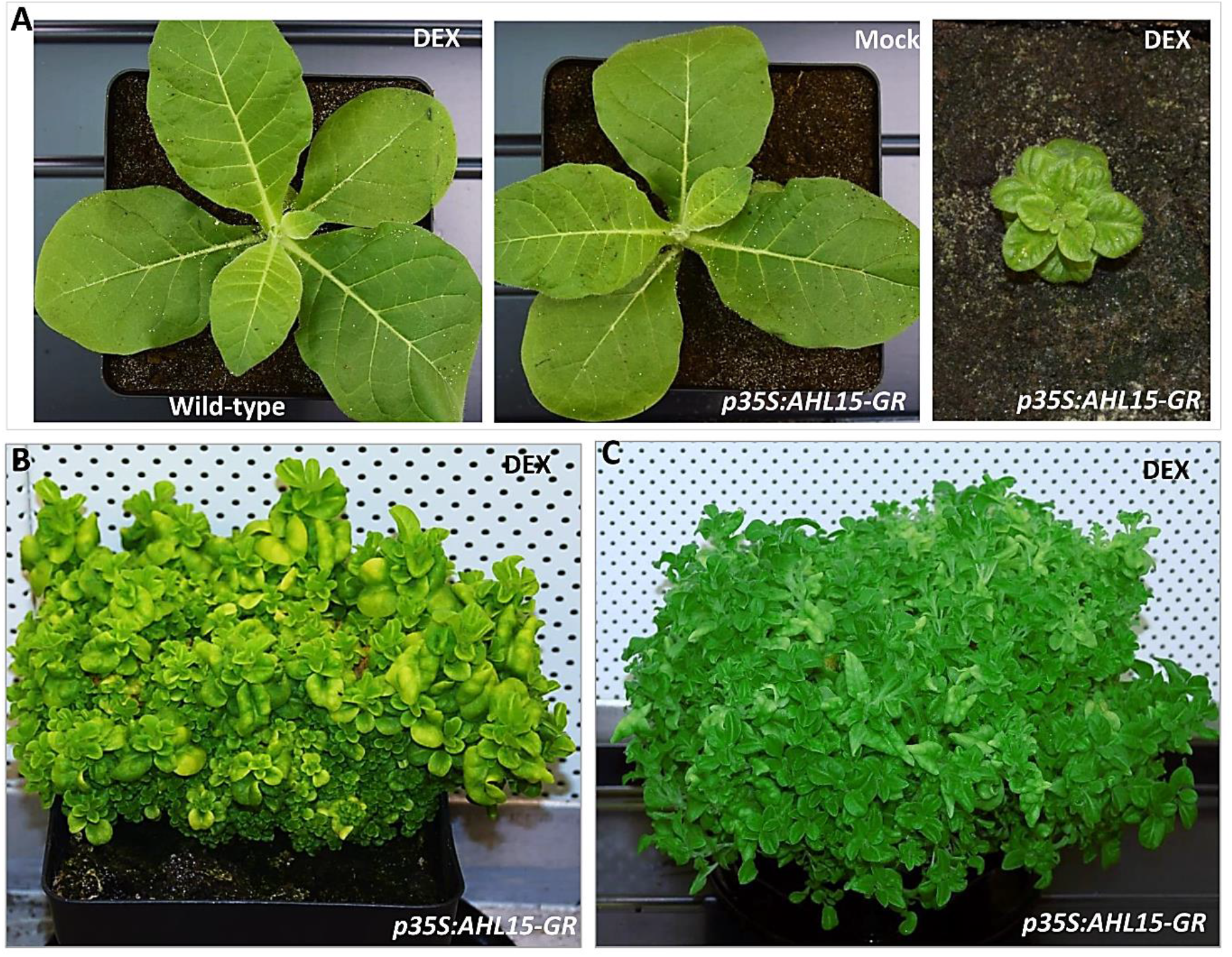
Extreme delay of the VPC by heterologous *AHL15* expression in tobacco. (**A**) Shoot morphology of a one-month-old wild-type (left) or *p35S:AHL15-GR* (middle and right) plant sprayed with water (Mock, middle) or sprayed with 20 μM DEX (DEX, left and right). (**B**, **C**) Extreme delay of the VPC in a six-month-old (**B**) or a one-year-old *p35S:AHL15-GR* (**C**) plant sprayed every week with 20 μM DEX.

**Supplementary Fig. 3.**
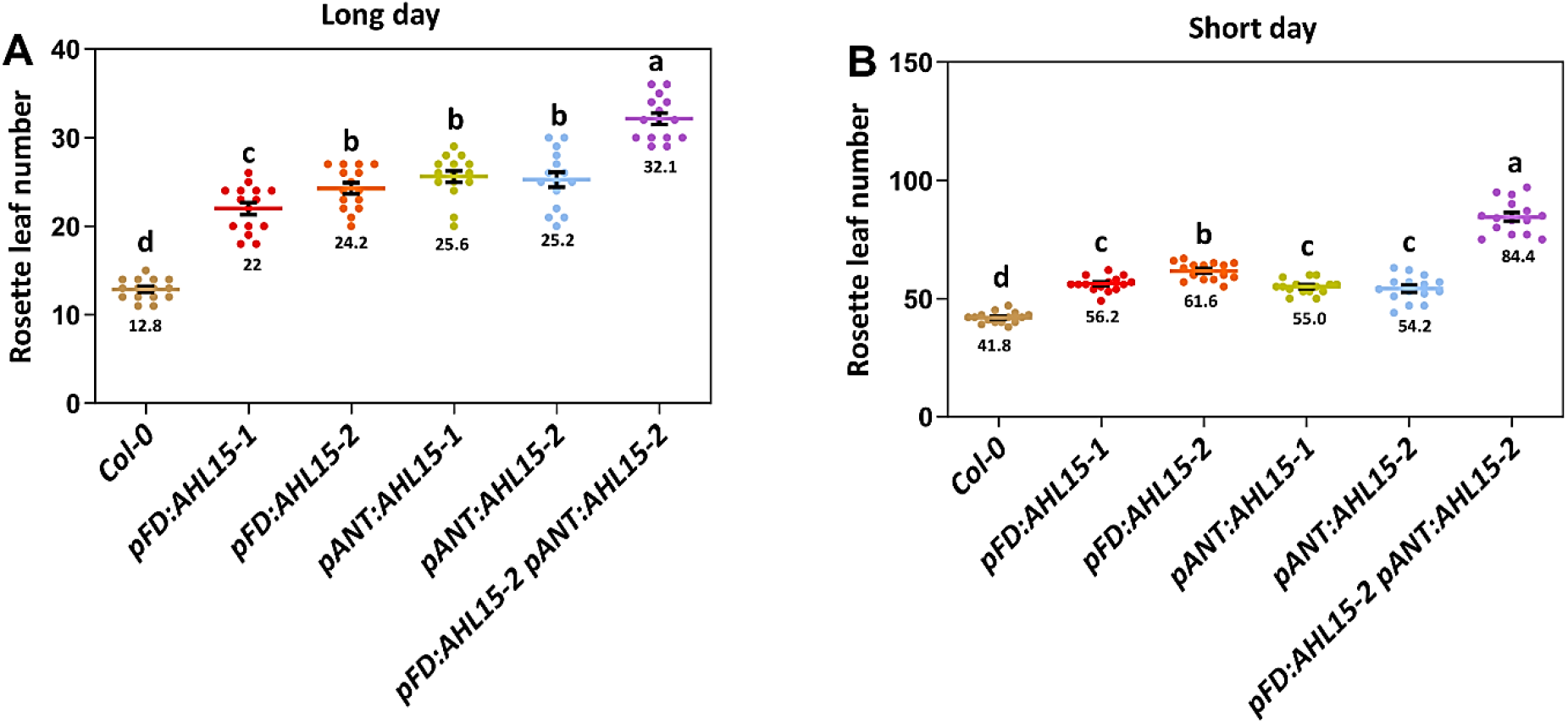
The effect of SAM- and young leaf-specific *AHL15* overexpression on flowering time. (**A**, **B**) The number of rosette leaves produced until flowering by wild-type (Col-0), *pFD:AHL15*, *pANT:AHL15* and *pFD:AHL15 pANT:AHL15* plants grown under LD (**A**) or SD (**B**) conditions. Colored dots indicate the number of rosette leaves per plant (n = 15 biologically independent plants) per line, horizontal line and the number below this line indicate the mean and error bars indicate the s.e.m. Different letters indicate statistically significant differences (P < 0.01) as determined by a one-way ANOVA with Tukey’s honest significant difference post hoc test.

**Supplementary Fig. 4.**
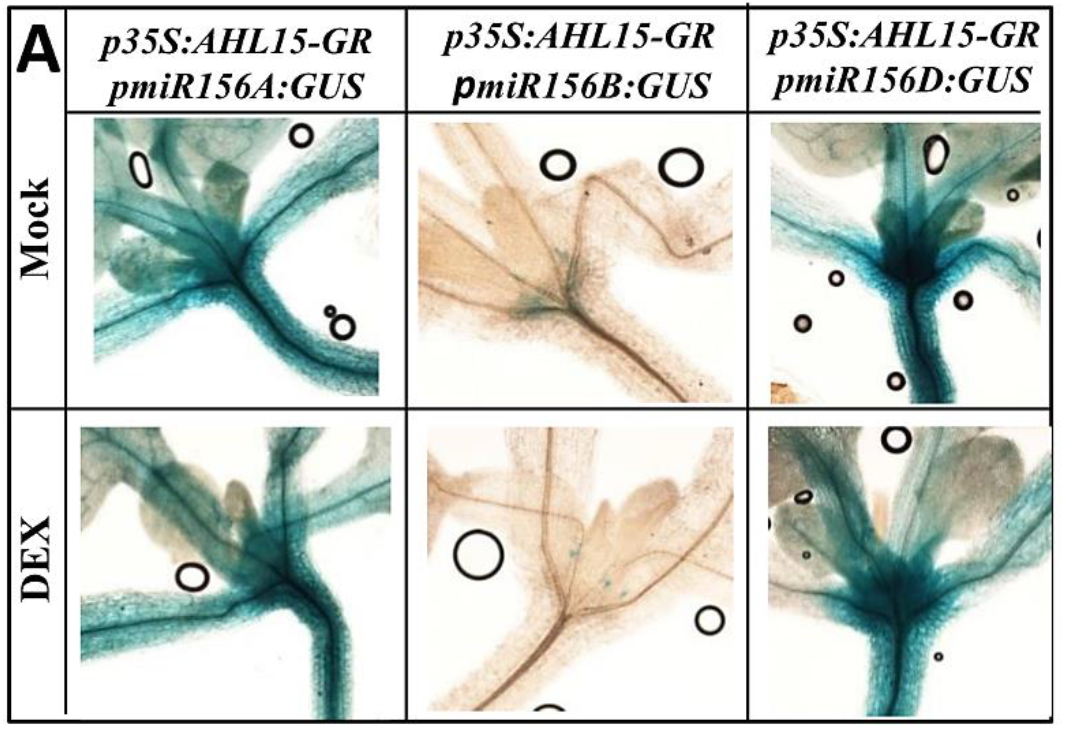
*AHL15* does not affect the expression of *miR156A,B,D*. **(A)** Histochemical staining for GUS activity in 2-week-old seedlings transgenic for *p35S:AHL15-GR* and *pmiR156A:GUS, pmiR156B:GUS* or *pmiR156C:GUS* following treatment with water (Mock, top) or DEX (bottom).

**Supplementary Fig. 5.**
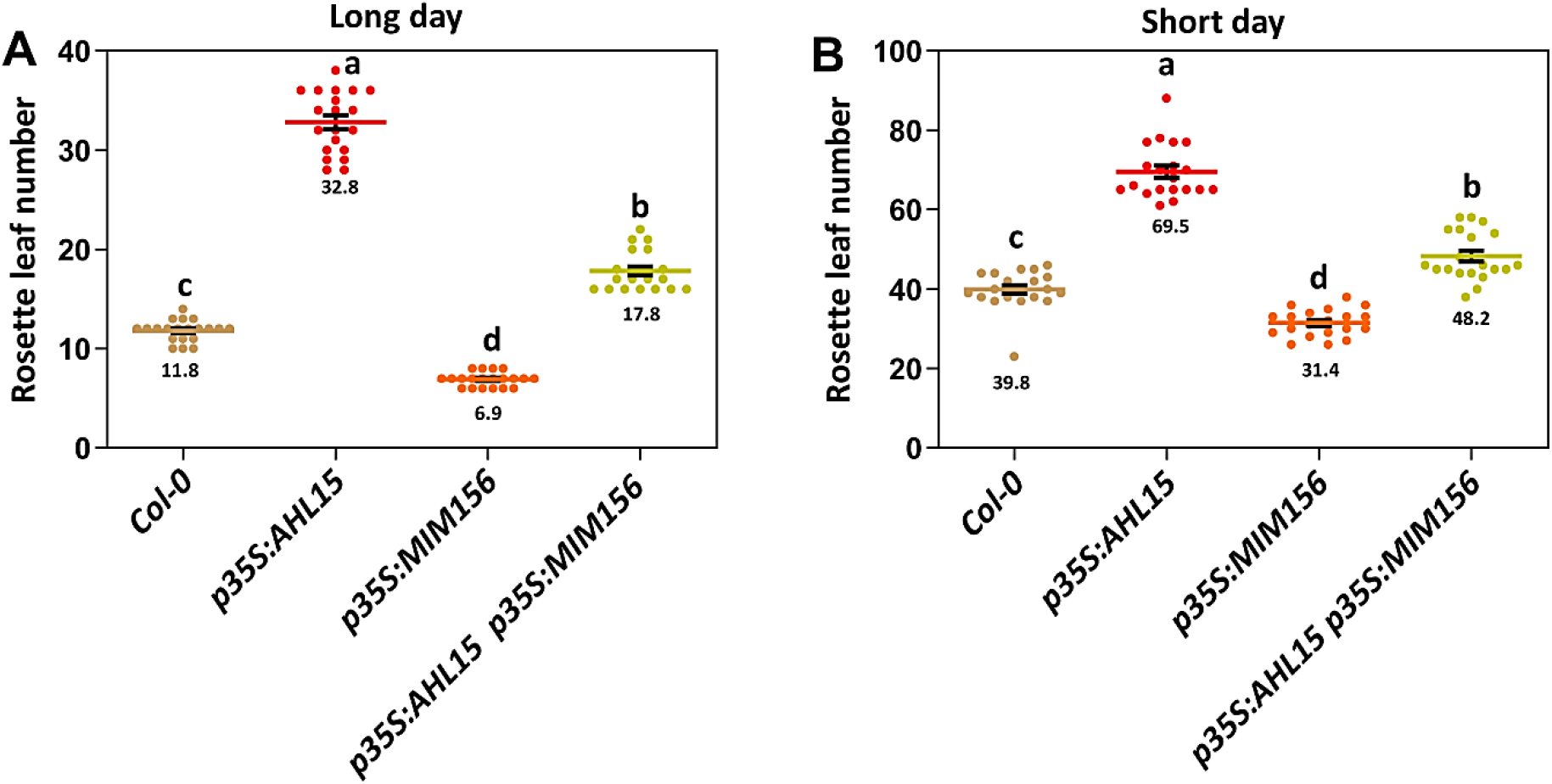
AHL15 and SPLs antagonistically control flowering time. (**A**, **B**) The number of rosette leaves produced until flowering by wild-type (Col-0), *p35S:AHL15*, *p35S:MIM156* and *p35S:AHL15*, *p35S:MIM156* plants grown under LD (**A**) or under SD (**B**) conditions. Colored dots indicate the number of rosette leaves per plant (n = 15 biologically independent plants) per line, horizontal line and the number below this line indicate the mean and error bars indicate the s.e.m. Different letters indicate statistically significant differences (P < 0.01) as determined by a one-way ANOVA with Tukey’s honest significant difference post hoc test.

**Supplementary Fig. 6.**
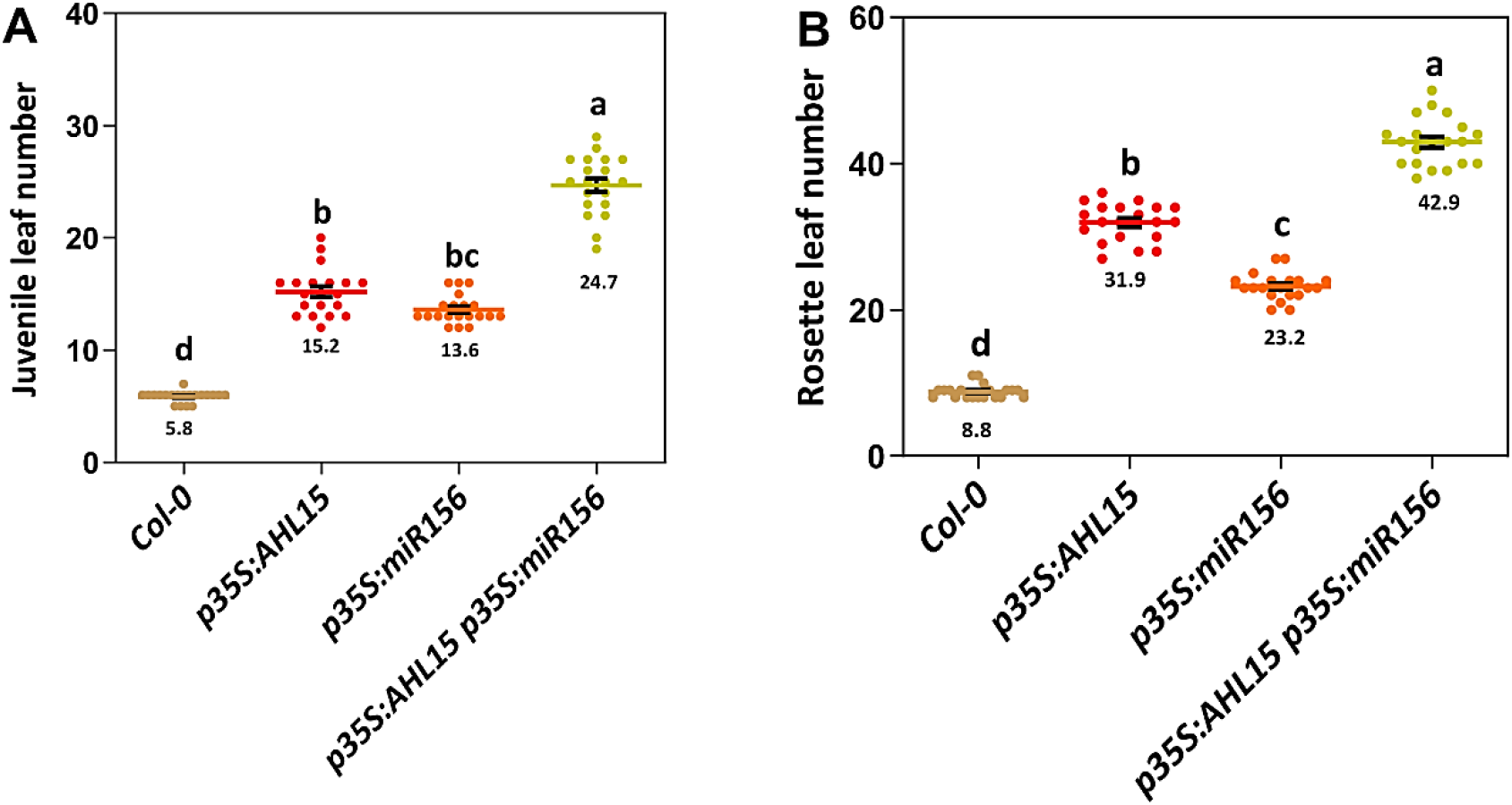
AHL15 and SPLs control the VPC and flowering time through parallel pathways. (**A**, **B**) The juvenile leaf number (leaves without abaxial trichomes) (**A**) and the number of rosette leaves produced until flowering (**B**) in wild-type, *p35S:AHL15*, *p35S:miR156* and *p35S:AHL15 p35S:miR156* plants grown under LD conditions. Colored dots indicate the number of leaves without abaxial trichomes (**A**) and the number of rosette leaves (**B**) per plant (n = 15 biologically independent plants) per line, horizontal line and the number below this line indicate the mean and error bars indicate the s.e.m. Different letters indicate statistically significant differences (P < 0.01) as determined by a one-way ANOVA with Tukey’s honest significant difference post hoc test.

**Supplementary Fig. 7.**
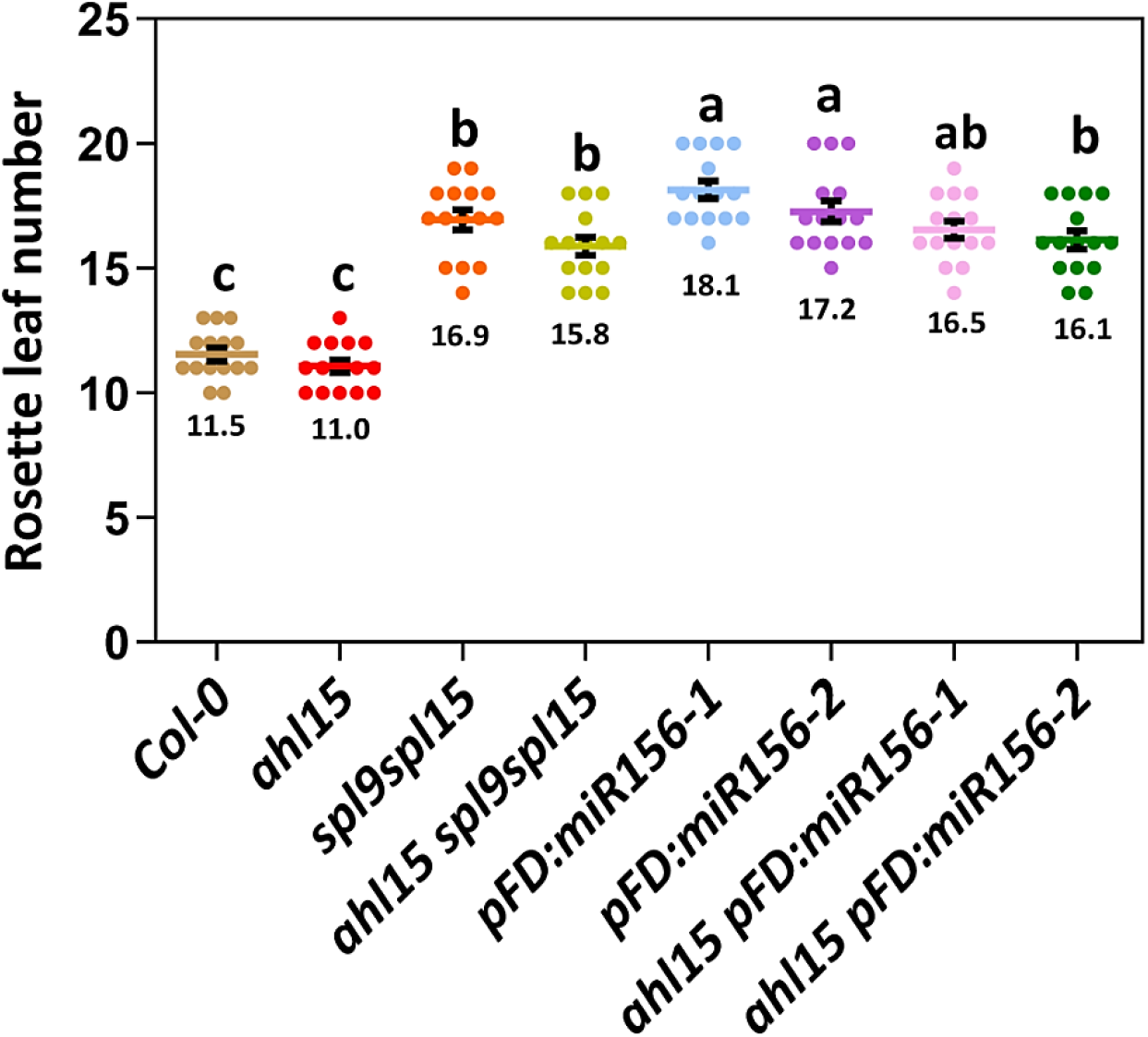
Delay of flowering by *spl* loss-of-function is largely *AHL15*-independent. The number of rosette leaves produced until flowering by wild-type (Col-0), *ahl15*, *spl9 spl15*, *ahl15 spl9 spl15, pFD:miR156*, *ahl15 pFD:miR156* plants grown under LD conditions. Colored dots indicate the number of rosette leaves per plant (n = 15 biologically independent plants) per line, horizontal line and the number below this line indicate the mean and error bars indicate the s.e.m. Different letters indicate statistically significant differences (P < 0.01) as determined by a one-way ANOVA with Tukey’s honest significant difference post hoc test.

**Supplementary Fig. 8.**
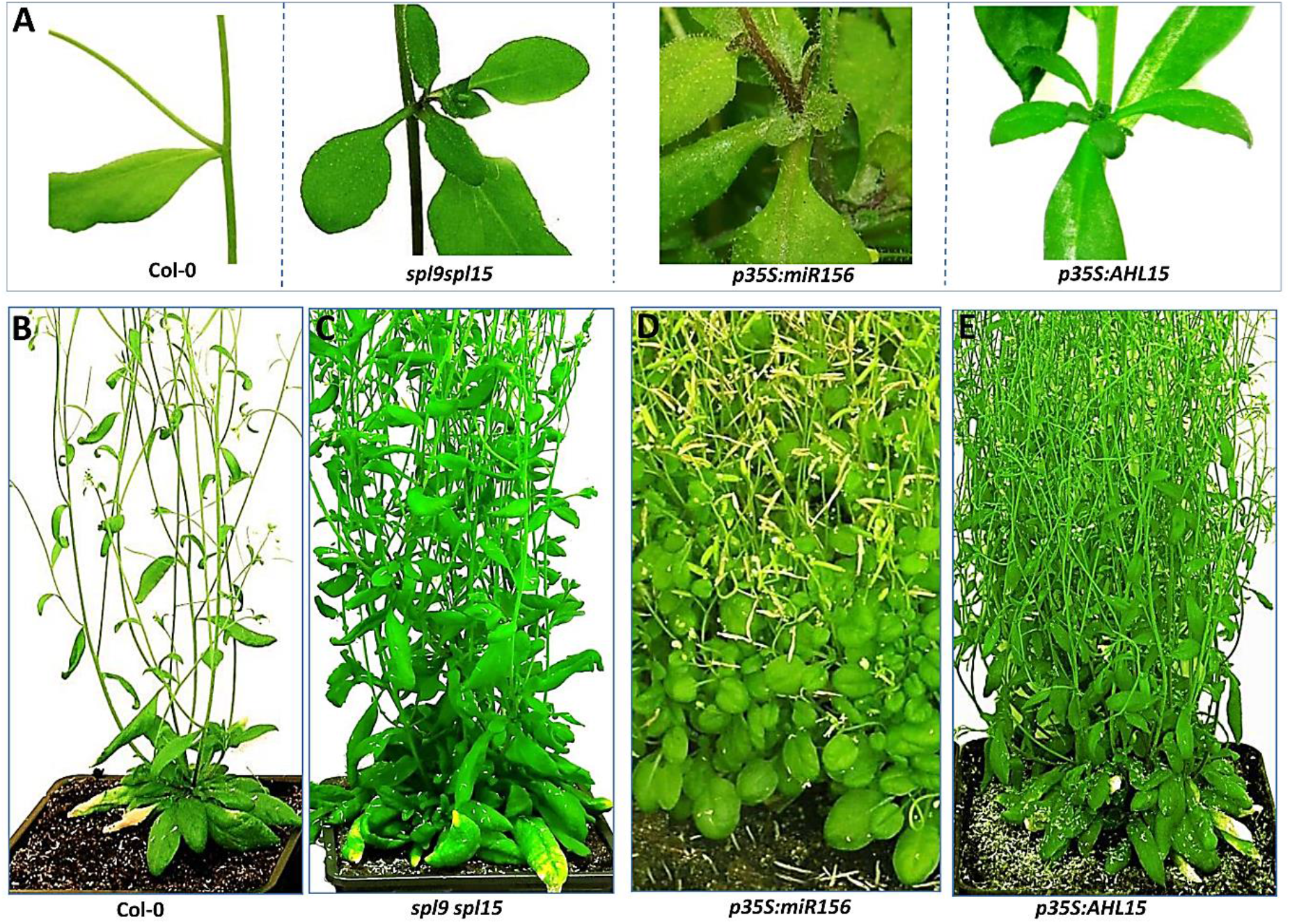
Aerial rosettes in Arabidopsis by reduced *SPL* expression or *AHL15* overexpression. (**A**,**B**) A detail of the first node of an inflorescence (B) and the complete shoot phenotype of a flowering wild-type (Col-0), *spl9 spl15*, *p35S:miR156* or *p35S:AHL15* plant grown under LD conditions.

**Supplementary Fig. 9.**
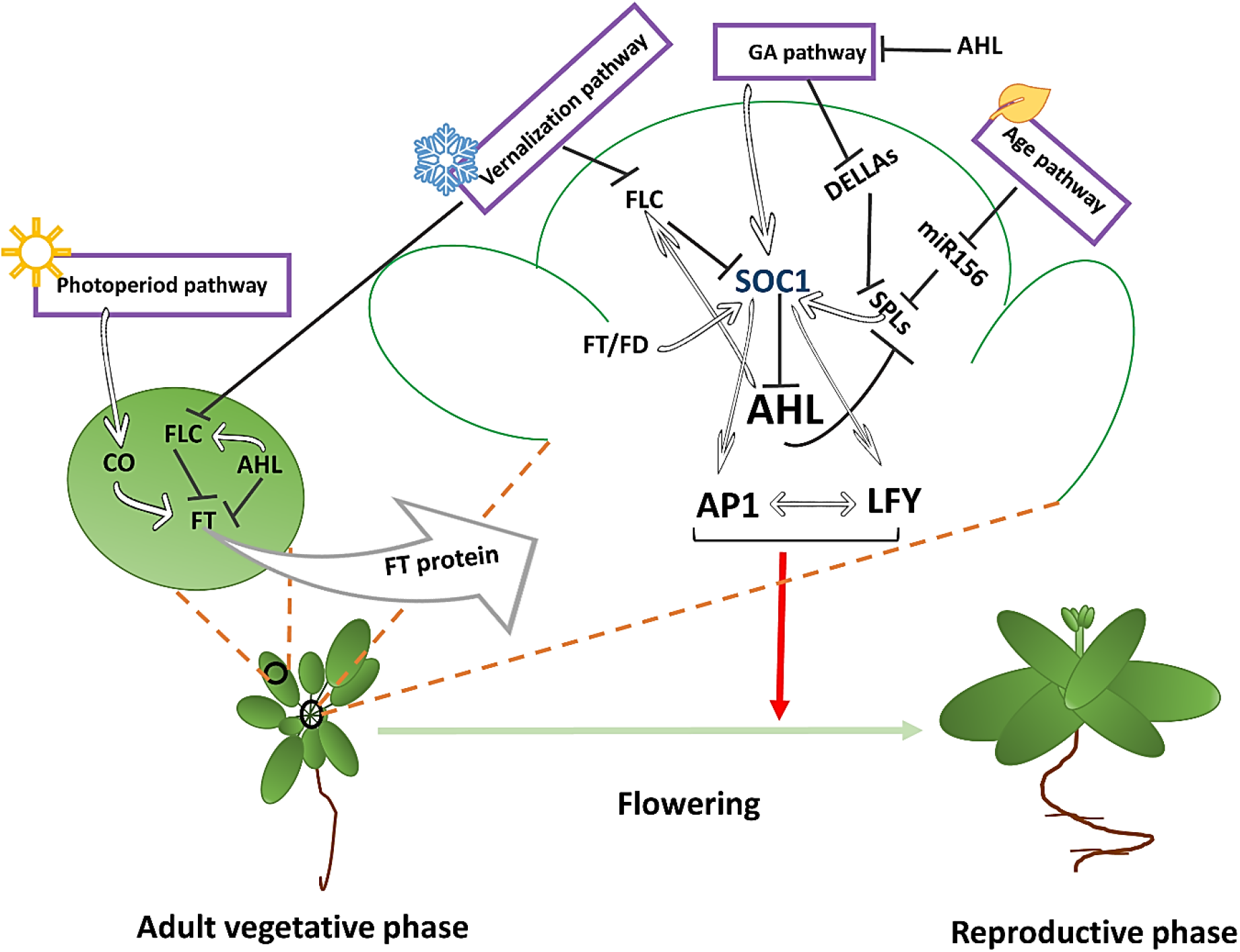
The position of AHLs in the network of pathways controlling flowering in Arabidopsis. In the vernalization pathway, cold treatment leads to stable repression of *FLC* transcription. The MADS box protein FLC determines the cold-period-dependent timing of flowering in Arabidopsis by repressing the expression of the floral integrator genes *FT* and *SOC1*. *FT* expression is induced in leaves by the photoperiod pathway through the accumulation of CO under long days. The FT protein subsequently travels to the SAM, where it physically interacts with FD to activate *SOC1*, which subsequently activates *AP1* and *LFY* (Song et al., 2013).AHLs are integrated into the vernalization pathway and the photoperiod pathway by activating FLC or directly repressing *FT* expression (Yun et al., 2012). In the age pathway, an age-dependent decline in miR156/miR157 levels allows an increased production of the SPL transcription factors, which activate the transcription of *SOC1* and other floral integrators (not shown). AHLs are integrated into the age pathway by repression of *SPL* expression. The phytohormone GA independently promotes flowering through activation of *SOC1* (and *SPL* expression). AHLs by repression of GA biosynthesis are integrated into the GA pathway. The subsequent activation of the downstream floral meristem identity genes, such as *LFY* and *AP1*, completes the floral transition.

**Supplementary Table 1.**
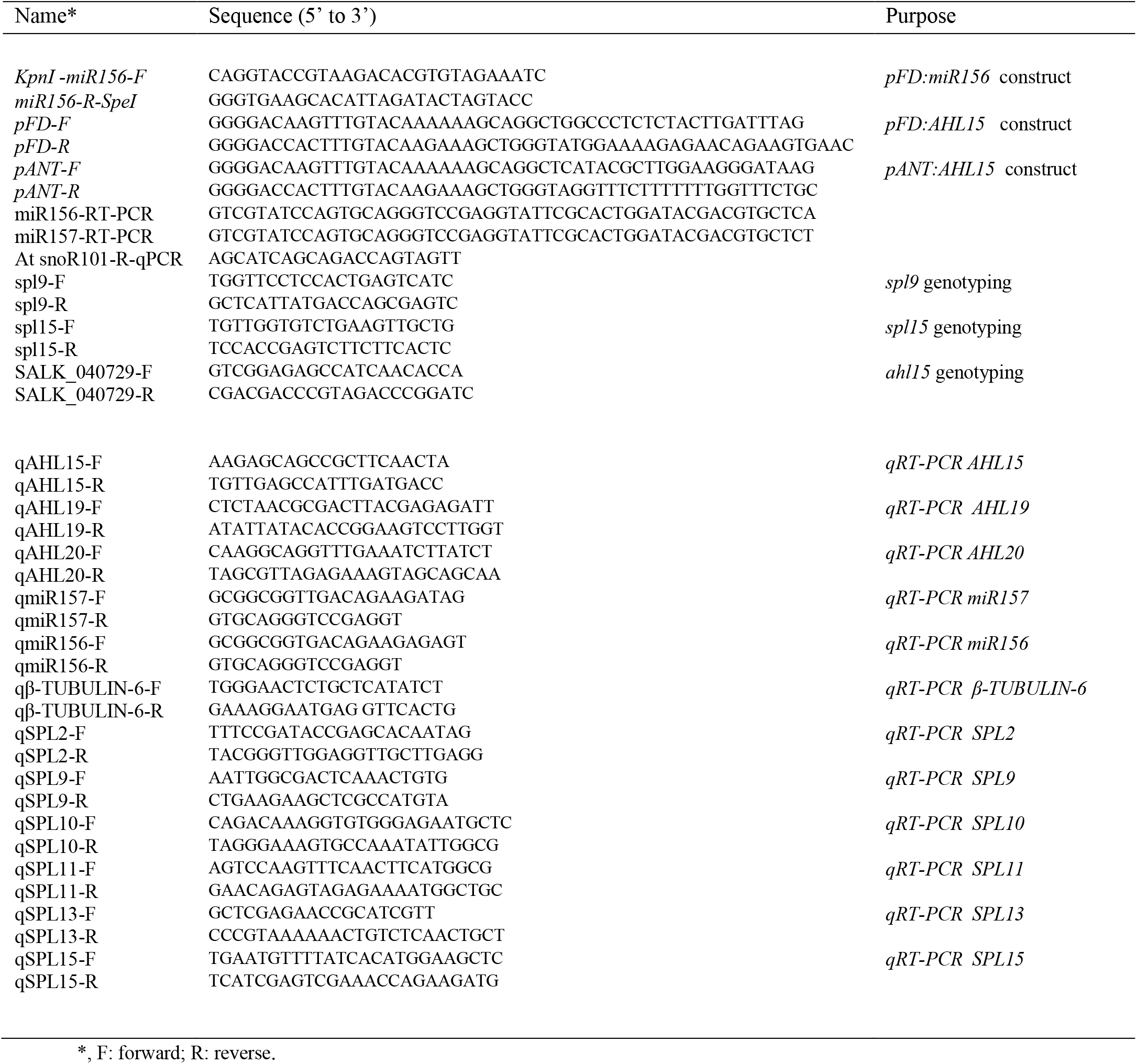
Primers used for cloning, genotyping and qRT-PCR

